# From Synthetic to Biological Nitrification Inhibition: Advancing Stabilization of Organic Fertilizers

**DOI:** 10.1101/2025.05.29.656819

**Authors:** Izargi Vega-Mas, Aude Mancia, Lucas Maggetto, Hugo Murillo, Alain Debaq, Bernard Heinesch, Francois Boland, Hans-Martin Krause, Hervé Vanderschuren, Cécile Thonar

**Author notes:** BioEcoAgro Joint Research Unit, INRAE, 02000, Barenton-Bugny, France.

## Abstract

Fertilizer type plays a critical role in nitrogen (N) cycling, influencing nitrous oxide (N_2_O) emissions, soil mineral N dynamics, and microbial communities. Understanding these interactions is essential for developing sustainable fertilization strategies that balance agricultural productivity with environmental protection. This study examined the effects of mineral and organic fertilizers (OFs) on N transformations and evaluated the efficiency of the nitrification inhibitor 3,4-dimethylpyrazole phosphate (DMPP) in mitigating N_2_O losses. Results showed that OFs exhibited variable impacts on N_2_O emissions depending on their composition and C/N ratio. DMPP effectively reduced nitrification-driven N_2_O emissions, particularly in treatments with high ammoniacal N content. However, its efficiency was limited with animal-based OFs, suggesting a complex interaction between fertilizer properties and inhibitor effectiveness. DMPP had not direct impact on soil microbial diversity but specifically targeted the *Nitrosomonaceae* family and *Nitrospira* class. Beyond synthetic inhibitors, biological nitrification inhibition (BNI) emerged as a promising alternative, which we explored using rhizospheric soils from wheat landrace Persia 44 and white mustard (cv. Pole Position, Verdi). These soils significantly reduced N_2_O emissions, particularly when combined with OFs. The integration of BNI with organic fertilizers, especially liquid digestate, represents a promising strategy for reducing N losses while maintaining soil fertility. This study underscores the need for tailored fertilization strategies that combine chemical and biological tools to optimize N use efficiency and support environmentally sustainable agriculture.

## 1. Introduction

Agriculture of the 21^st^ century faces critical challenges. Food production must intensify to accommodate the global population which is expected to reach nine billion by 2050 (United Nations, 2019). Simultaneously, agriculture must mitigate its negative impacts on the environment and human health. This sector currently contributes to approximately 30% of global greenhouse gases (GHG) emissions (Lynch et al., 2021). Moreover, it significantly contributes to important nitrogen (N) losses, including nitrous oxide (N_2_O) emission, ammonia (NH_3_) volatilization, and nitrate (NO_3_ ^−^) leaching. While N is essential for crop production, about 50-70 % of N inputs are lost due to the poor N use efficiency (NUE) of agroecosystems (Cassman et al., 2002). This issue stems from an unbalanced N cycle, characterized by the accumulation of reactive N within agroecosystems (Erisman et al., 2013). Therefore, it is urgent to develop new technologies and practices that can reduce GHG emissions and mitigate N losses from agricultural systems.

The use of nitrification inhibitors (NIs) represents an interesting strategy to increase NUE of agroecosystems and to reduce N losses by developing low soil nitrifying conditions. Nitrification is a crucial step in the N cycle, comprising the aerobic oxidation cascade from NH_4_ ^+^ to nitrate NO_3_ ^−^, via nitrite (NO_2_ ^−^) and hydroxylamine as intermediates (Ward, 2015). During this process, polluting gases such as nitric oxide (NO) and N_2_O can be emitted (Pilegaard, 2013). Additionally, the NO_3_ ^−^ produced may be lost through leaching or denitrification, the later also contributing to NO and N_2_O emissions (Saggar et al., 2013). Nitrification involves autotrophic microorganisms, including, archaea and bacteria such as ammonium oxidizing bacteria/archaea (AOB and AOA, respectively) and nitrite oxidizing bacteria (NOB) (Ward, 2015).

Common synthetic nitrification inhibitors (SNIs) like dicyandiamide (DCD) and 3,4-dimethylpyrazole phosphate (DMPP) inhibit the enzyme ammonia monooxygenase (AMO), which catalyzes the first step of nitrification (Benckiser et al., 2013; Chaves et al., 2006). By extending the retention time of NH_4_^+^ in the soil, these inhibitors improve NUE in crops and reduce N losses (Cameron et al. 2013; Cookson & Cornforth, 2002). Meta-analyses have shown that DCD and DMPP can reduce N_2_O emissions by 40-50% (Gilsanz et al., 2016; Yang et al., 2016a; Lei et al., 2022). However, the efficiency of NIs depends on soil properties such as soil texture and pH, which vary significantly between studies (Guo et al., 2023; Tufail et al., 2023), and onfertilizer type (Peixoto and Petersen, 2023; Chen et al., 2013; Lazcano et al., 2021). Most meta-analyses classify fertilizers into chemical or organic categories, rarely distinguishing among different types of organic fertilizers (OFs). Nevertheless, it is important to assess the efficiency of NIs across diverse OFs, which offer benefits to the soil compared to chemical fertilizers, such as carbon (C) sequestration (Yan & Gong, 2010), increased microbial activity and enhanced nutrient cycling (Lazcano et al., 2013). Furthermore, incorporating OFs can help decrease fossil fuel consumption associated with synthetic fertilizer production, thereby supporting efforts to mitigate GHG emissions. Additionally, the revalorization of agricultural residues contributes to not only aids in nutrient recovery but also reinforces the principles of a circular economy (Haque et al., 2023).

In order to advance towards environmentally friendly agriculture without compromising crop yields, it is essential to develop new and sustainable technologies for controlling soil nitrification, ensuring that the benefits provided by nitrification inhibitors (NIs) are maintained. Over the last two decades, research has increasingly focused on nature-based solutions, such as biological nitrification inhibition (BNI) (Subbarao and Searchinger, 2021; Wang et al., 2021; Saud et al., 2022). Biological nitrification inhibitors (BNIs) are organic compounds produced by certain plants in their rhizosphere and/or in tissues to suppress soil microbial nitrification. This trait was firstly identified in the tropical grass *Brachiaria humidicola* and in *Sorghum bicolor* (Subbarao et al., 2007) and has since been described in a larger number of species including *Brassicaceae* (Brown and Morra, 2009), or cereals such as rice and maize (Otaka et al., 2021; Wang et al., 2021). Although current elite wheat varieties show weak BNI capacity (Subbarao et al., 2007b), high BNI potential has been identified in a set of wheat landraces (O’Sullivan et al., 2016). In the context of practical application, BNIs offer several advantages over SNIs, including lower costs for farmers, supposedly reduced environmental impact, continuous production by plants, and greater social acceptance (Coskun et al., 2017; Sadhukhan et al., 2022). Moreover, using BNI-producing plants to stabilize the ammonium fraction of OFs could enable the development of mitigation strategies compatible with organic farming systems.

To the best of our knowledge, no study has yet directly compared the efficiency of DMPP and BNI-containing soils in reducing nitrification and N losses across a range of organic fertilizers under controlled conditions. While most research on NIs and OFs has focused on N_2_O emissions, studies rarely integrate other critical nitrification parameters, such as changes in soil mineral N pools, or the abundance and composition of bacterial populations, including nitrifiers (Li et al., 2023; Severin et al., 2016; ten Huf & Olfs, 2020). In this study, we not only evaluated the impact of the synthetic nitrification inhibitor DMPP across different OFs but also advanced the field by exploring the stabilization potential of BNIs as a novel, biologically derived alternative to synthetic inhibitors, focusing on the two most promising fertilizers.

## 2. Materials and methods

### 2.1. Soil sampling and experimental setup

The first part of this study was based on a 48 days soil incubation experiment (EXP 1), with two types of samplings: a non-destructive gas sampling to measure N_2_O emissions and a destructive soil sampling to assess pH, mineral N, dissolved organic carbon (DOC), selected N cycle-related microbial genes abundance and bacterial composition by sequencing. For the purpose of this soil incubation experiment, a silt loam alkaline soil (10.3 % sand, 76.2 % slit, 13.5 % clay) was collected from the 0-10 cm layer of a former sugar beet field in the locality of Gembloux, Belgium (50°35’00.3“N 4°41’23.1“E). The soil was air-dried for four days and then homogenized, sieved at 3.5 mm and kept at 4 °C until use. Subsamples of the soil were sent to Laboratoires d’Analyses Agricoles, La Hulpe, Belgium for classical soil analyses (**Table S1**). For EXP1, the incubation microcosm was set up in 250 ml glass bottles filled with fresh soil equivalent of 200 g of dried soil, with four technical replicates per treatment and per destructive sampling time point (4 replicates x 11 treatments x 4 time points). Water was added to each pot up to WPFS value close to 35%; WFPS (Water-Filled Pore Space) is calculated as follows: WFPS = (soil gravimetric water content × soil bulk density) / (1 – (soil bulk density / particle density)) × 100, where soil bulk density was 1.01 g cm^−3^ and particle density was assumed to be 2.65 g cm^−3^. Pots were closed with pierced parafilm in order to maintain humidity while allowing gas diffusion and were covered with aluminum foil to ensure dark conditions. Pots were then placed in a controlled growth chamber at 21-24°C for a one-week acclimation phase before addition of fertilizers (= start of the experiment).

Fertilizers were applied at Day 0 on soil microcosms, at a rate of 100 mg N kg^−1^ DW soil. Fertilizer treatments were ammonium sulfate (AS); two solid organic fertilizers (OFs): Orgamine 7 (O7) which are commercial solid pellets of plant and animal origin, and chicken manure (CM) mixed with wheat straw; as well as two liquid OFs: liquid digestate (D) derived from the bio-methanization of vegetable wastes, and cattle slurry (CS); and a non-fertilized control (NF). Ammonium-N percentage (% of total Nitrogen) of the fertilizers were 100, 46, 31, 13 and 31% for AS, D, O7, CM and CS respectively. More information about fertilizers characteristics are provided in **Table S2**. The fertilizer treatments were also applied in combination with DMPP, at a rate of 1% of applied N, coming out as treatments AS+, O7+, CM+, D+ and CS+ respectively. Therefore, 11 treatments were assessed in this study (EXP 1). In order to achieve homogeneous distribution of fertilizers in the soil, mineral fertilizer (AS) was first dissolved in a small volume of deionized water and applied evenly on the soil surface area of the microcosms, same as for liquid organic fertilizers (AS, D and CS). Solid fertilizers (O7 and CM) were previously grinded and applied by mixing with the soil. Pots of all treatments of EXP 1 and EXP 2 were subjected to soil mixing for methodological homogeneity. All the pots were finally adjusted to 60 % WFPS and adjusted individually every 2-3 days until the end of the incubation.

An additional 21-day incubation experiment (EXP 2) was subsequently set up to compare the efficiency of the synthetic NI (DMPP) with that of rhizospheric soils collected from plant species with demonstrated and contrasted BNI (Biological Nitrification Inhibition) activity. A similar set-up to EXP 1 was established for EXP 2, with 200 g equivalent DW soil soil (mixed with sand at 2:1, w:w) packed in 250 ml glass bottles and treated according to the following treatment design. The first studied factor was the type of treatment previously applied to the soil, which included: not treated soil (NT), soil treated with DMPP (SDMPP), and soil collected from the rhizosphere of wheat cv. Persia 44 (P44), white mustard cv. Verdi (VERDI) and white mustard cv. Pole Position (PP) (see **supplementary material Text S1 and Table S1,** for the methodology and properties of the collected rhizospheric soil). The selected cultivars of wheat (*Triticum aestivum*) and white mustard (*Sinapis alba*) were chosen for their ability to reduce, in a contrasted way, nitrification rate in either hydroponic or soil-based studies (Brown and Morra, 2009; O’Sullivan et al., 2016; Jauregui et al., 2023). The second factor was the type of fertilizer supplied in the 21-day incubation phase. Fertilizer was applied at a rate of 100 mg N kg^−1^ DW soil as AS (ammonium sulfate), O7 (Orgamine 7), or D (liquid digestate), with four technical replicates per treatment and per destructive sampling time point (4 replicates x 5 soil treatments x 3 fertilizer types x 4 time points). Four replicates of a negative control (NC) consisting of untreated and unfertilized soil were also included for each time point.

### 2.2. N_2_O measurements (EXP 1 and EXP 2)

Nitrous oxide (N_2_O) fluxes were measured following the dynamic chambers method as described in (Lognoul, 2019), which in this case consisted in hermetic closed glass pots connected to a hermetic circuit in which the air was circulating to the gas analyser. For EXP 1, N_2_O fluxes were first determined daily, starting from 4 h after the application of fertilization and up to Day 10, then each two days up to Day 22, and finally each three days from Day 28 to 48. The measurements were performed with a calibrated Thermo ScientificTM 46i infra-red analyser, Waltham, USA). Nitrous oxide (N_2_O) concentration (in ppm) was measured at regular intervals (30 seconds) during the entire enclosure time of 15 minutes. Then the chamber opened again for 90 seconds to purge the system, and the next pot was closed and measured. The measurements were performed automatically in four series of 11 treatments plus and empty pot (as control of the background signal). N_2_O emission was then converted in µg N-N_2_O units per g of dry soil and hour, using the 15-minute slope value, and correcting this value with the pot headspace volume and N_2_O water solubility. Cumulative N_2_O emissions were calculated by linear interpolation of daily fluxes.

For EXP 2, N_2_O fluxes were determined daily during the first week and then every 2-3 days until the end of the incubation (i.e. 21 days), starting from few minutes after fertilizer application (i.e. Day 0). Fluxes were measured with the Li-7820 N_2_O/H_2_O (LICOR ®). For this, an adapted lid of the 250mL glass pots was drilled with two holes allowing the introduction of the tubing of a closed-loop system attached to the gas analyzer. Contrary to EXP 1, the measurements were made manually. Therefore, after closure of one pot with the previously described lid, gas sample was collected during 3 min and the lid was then removed and inserted onto the following pot for another 3 min measurements. The N_2_O flux was calculated according to the 3 min slope, and cumulative emissions were calculated by linear interpolation of daily fluxes as in EXP 1.

### 2.3. Destructive soil samplings and analyses (EXP 1 and EXP2)

For EXP1, immediately after fertilizers application, a batch of soil pots was destructively sampled for determination of physicochemical and microbial characteristics at Day 0. Subsamples of each treatment were also sent to Laboratoires d’Analyses Agricoles, La Hulpe, Belgium to determine total N and C contents of the soil-fertilizer mixtures (**Fig. S1**). Other destructive samplings occurred at Day 7, Day 15 and Day 48 of the soil incubation experiment.

Change in soil mineral N (ammonium NH_4_ ^+^-N, nitrate NO_3_ ^−^-N and nitrite NO_2_ ^−^-N) and dissolved organic carbon (DOC) contents were determined in the four replicates per treatment at each destructive sampling time point. Soil of each pot was collected and thoroughly homogenized. In order to determine soil NH_4_ ^+^-N, NO_3_ ^−^-N and NO_2_ ^−^-N, 50 g of fresh soil were mixed with 1 M KCl (1:2, w:v) and shaken for 1 h at 165 rpm. The soil solution was settled for 1 hour, and then filtered through Whatman no. 1 filter paper (GE Healthcare, Little Chalfont, Buckinghamshire, UK). One fraction was further filtered through a Sep-Pak Classic C18 Cartridge (125 Å pore size; Waters, Milford, MA, USA) to eliminate organic carbon. Mineral N contents were determined spectrophotometrically in 96-well plates (Tekan microplate reader, Spark 10K, Switzerland). Concentration of NH_4_ ^+^-N was determined with the Berthelot method (Patton & Crouch, 1977) while the concentration of NO ^—^Nwas determined by the VCl_3_-Griess method (Miranda et al., 2001), in which NO_3_ ^−^ is reduced to NO_2_ ^−^ with VCl_3_, and thus the content of total NO_x_ is determined. Afterwards, the content of NO_2_ ^−^-N was determined with Griess reagents without using VCl_3_, and this value was subtracted to the total NO_x_ to deduce NO_3_ ^−^-N content. DOC contents were determined from the remaining fraction of the paper-filtered KCl extracts. Samples were further filtered with Sterile PES syringe filters - 0.45 μm (ROCC), and diluted to 1/5 and measured following the procedures regulated by ISO 10694:1995 at the Bureau Environment et Analyses (BEAGx), Gembloux Agro-Bio Tech, ULiege. Soil pH was determined in the four replicates per treatment at each time point. For this, 15 g of fresh soil were first oven-dried at 35°C. Dry samples were then suspended in deionized water (1:2.5, w:v) and shaken for 1 h at 165 rpm. Soil suspensions were then settled by centrifugation and pH was determined with a probe in the solution phase. Final value of each sample was taken with the pH probe after three minutes to assure the stabilization of the measurement.

For EXP 2, soil samples were taken first from the plant growing pots (as well as from non-treated pots), and then from incubations pots, at Day 0, 7 and 21 after the addition of the fertilizer. Mineral N was extracted by KCl as previously described. Concentration of NH_4_ ^+^-N and NO_3_ ^−^-N were determined with the same methods as for EXP 1.

### 2.4. DNA extraction and quantification of nitrifying and denitrifying gene abundance (EXP 1)

Bacterial and archaeal nitrification marker gene (*amoA*) for ammonium oxidizing bacteria (AOB) and ammonium oxidizing archaea (AOA), 16s rRNA gene for total bacterial and archaeal abundance, and denitrification bacterial marker genes of *narG, nirS, nirK, nosZI* and *nosZII* were quantified by quantitative PCR (qPCR) using soil DNA extracts as templates. For DNA extraction, soil subsamples were collected from the mixed soil microcosms at Day 0, 7, 15 and 48, then immediately frozen in liquid nitrogen and stored at −80 °C until use. DNA was then extracted from the equivalent of 0.25 g of dry soil using the DNeasy PowerLyzer PowerSoil Kit (Qiagen) and a Tissue Lyser (Retsch MM300, Qiagen) with Eppendorf adapters. Sample DNA concentration was quantified using QuantiFluor® ONE dsDNA System (Promega) in the QUANTUS fluorimeter following the manufacturer’s instructions. Marker genes were amplified by qPCR using KAPA SYBR FAST qPCR Universal Master Mix (Kapa Biosystems, Wilmington, MA) and a CFX96 Touch™ Real-Time PCR Detection System (Bio-Rad, Hercules, CA, USA). Each reaction was performed in 10 μl containing 1 μl of DNA template (diluted in a range of 10 ng per μl) and 9 μl of reaction mixture (see **Tables S2 and S3** for primer sequences, qPCR conditions and reaction compositions). Standard curves were prepared from serial dilutions of linearized plasmids (**Table S4**) with target gene insertions ranging from 10^8^ to 10^2^ gene copies μl^−1^. The copy number of the target gene per gram of dry soil was then quantified using the following equation, normalized for DNA concentration: [(number of target gene copies per reaction × volume of DNA extracted) / (volume of DNA used per reaction × gram of dry soil extracted)] / DNA concentration

### 2.5. DNA sequencing (EXP 1)

PCR amplification was performed in a total volume of 25 μl reaction mixture containing 10-40 ng of DNA template (10 µl of 1/30 diluted soil DNA extracts), 12.5 µl Q5 High-Fidelity 2X Master Mix (NEB M0492S) and 1.25 µl of each primer at 10 µM. The PCR conditions consisted of an initial denaturation at 98°C for 3 min, 30 amplification cycles of [95°C for 10 s, 58°C for 10 s and 72°C for 20 s], followed by a final elongation at 72°C for 2 min. Each sample was amplified in triplicates and pooled prior to purification with Agencourt AMPure XP beads (Beckman Coulter, Berea, CA) and quantification with the QuantiFluor® ONE dsDNA System (Promega) in the QUANTUS fluorimeter. Amplicon pools ranging for a final concentration of 10-18 ng DNA/µl were sent to the Génome Québec Innovation Center at McGill University (Montréal, Canada) for barcoding using the Fluidigm’s Access ArrayTM technology (Fluidigm) and paired-end sequencing on the Illumina MiSeq v3 (PE300) platform (Illumina Inc., San Diego, CA, USA).

### 2.6. Bioinformatic analysis (EXP 1)

A total of 7 837,983 raw paired end reads were obtained and processed using the Euler Scientific Compute Cluster at ETH Zürich. For most steps, USEARCH v11.0.667 (Edgar, 2010) commands were used with default settings unless stated otherwise. The positive control sequence of bacteriophage phi X and low-complexity reads were removed with filter_phix and filter_lowc, respectively. The remaining reads were trimmed with fastx_truncate and merged with fastq_mergepairs (min overlap 30, min %identity:60 and min merged length 100). Primer sequences were removed with search_pcr with two mismatched allowed. Reads were quality filtered, using PRINSEQ-lite v0.20.4 (Schmieder, 2011) with an allowed amplicon size range of 350-480 bp. Subsequently, UPARSE was employed to construct amplicon sequence variants (zero OTUs) and UNOISE3 was employed for denoising before an additional clustering step at 97% sequence identity via UPARSE-OTU (Edgar, 2013). The taxonomy for 16S rRNA gene data was assigned using SINTAX (Edgar, 2016) (sintax_cutoff 0.85) based on SILVA gene database (Quast et al.. 2013). Before downstream analysis non-bacterial (archaea, mitochondria, and chloroplasts) were removed, resulting in 3,874,789 bacterial sequences (34,608 to 50,457 sequences per sample) assigned to 6290 zOTUs (**Table S6)**.

### 2.7. Statistical analyses

Data were subjected to statistical analysis using IBM SPSS 21.0 statistical package (IBM, 2012) for EXP 1 and the software R studio version 3.2.5 (R Development Core Team, 2019) for EXP 2. Student-T test (p<0.05) or one-way ANOVA followed by Duncan’s test (p<0.05) were employed. Details of statistical analyses are given within Figure legends.

Community structure of soil bacteria was assessed using the phyloseq package (McMurdie and Holmes, 2013). ZOTUs with fewer than 10 reads occurring in less than 10% of samples were removed to reduce sparsity of microbiome data (Cao, 2021). Rarefaction plots were created via ggrare (**Fig. S2**). Prevalence filtered data was clr-transformed and a distance matrix based on euclidean distances was calculated before differences in community structure were assessed by Permanova, with 10^4^ permutations and time, fertilizer type and DMPP as experimental factors. Pairwise comparison between treatments was conducted using pairwise.adonis2 with p-value adjustment according to Benjamini–Hochberg. To target potential nitrifiers subsets for the family of *Nitrosomonadaceae* and the class *Nitrospirae* as well as the two timepoints of sampling were subjected to the same analysis (Table 1). To visualize differences of bacterial community composition the vegan package (Okasanen et al., 2012) was used for unconstrained ordination via NMDS and constrained ordination via dbRDA (Figure 4). To identify bacterial taxa that are sensitive to DMPP application, analysis of indicator species was performed using the multipatt function of the indicator species package (De Cáceres, 2012) with 10^4^ permutations, as described in Lori et al., 2022.

**Table 1.** Effect of fertilization, incubation time and DMPP amendment on bacterial community structure and OTUs assigned to family of *Nitrosomonadaceae* and class of *Nitrospira*, as assessed by permutational analysis of variance (PERMANOVA). Bold letters represent p values < 0.05.

|  | Df | Total bacteria |  |  | Family *nitrosomonas* |  |  | Class *Nitrospirae* |  |  |
| --- | --- | --- | --- | --- | --- | --- | --- | --- | --- | --- |
|  |  | F. Model | R2 | Pr(>F) | F. Model | R2 | Pr(>F) | F. Model | R2 | Pr(>F) |
| fert | 5 | 2.896 | 0.14 | **>0.001** | 1.342 | 0.07 | **0.003** | 1.815 | 0.1 | **0.005** |
| time | 1 | 5.445 | 0.05 | **>0.001** | 2.739 | 0.03 | **>0.001** | 1.191 | 0.01 | 0.286 |
| dmpp | 1 | 1.026 | 0.01 | 0.32 | 2.072 | 0.02 | **0.004** | 1.258 | 0.01 | 0.258 |
| fert:time | 5 | 1.556 | 0.07 | **>0.001** | 1.186 | 0.06 | <u>0.053</u> | 1.2 | 0.06 | 0.199 |
| fert:dmpp | 4 | 1.014 | 0.04 | 0.364 | 1.001 | 0.04 | 0.478 | 0.941 | 0.04 | 0.557 |
| time:dmpp | 1 | 0.991 | 0.01 | 0.44 | 1.147 | 0.01 | 0.258 | 1.401 | 0.02 | 0.189 |
| **T7** |  |  |  |  |  |  |  |  |  |  |
| fert | 5 | 2.584 | 0.25 | **>0.001** | 1.352 | 0.15 | **0.004** | 1.773 | 0.19 | **0.007** |
| dmpp | 1 | 1.029 | 0.02 | 0.317 | 2.19 | 0.05 | **>0.001** | 0.439 | 0.01 | 0.892 |
| fert:dmpp | 4 | 1.009 | 0.08 | 0.407 | 1.087 | 0.09 | 0.239 | 1.073 | 0.09 | 0.358 |
| **T48** |  |  |  |  |  |  |  |  |  |  |
| fert | 5 | 1.897 | 0.2 | **>0.001** | 1.197 | 0.13 | <u>0.062</u> | 1.247 | 0.13 | 0.169 |
| dmpp | 1 | 0.989 | 0.02 | 0.45 | 0.984 | 0.02 | 0.495 | 2.495 | 0.05 | **0.018** |
| fert:dmpp | 4 | 1.005 | 0.08 | 0.416 | 1.144 | 0.1 | 0.143 | 1.391 | 0.12 | <u>0.091</u> |

## 3. Results

### 3.1. Soil N_2_O emissions in presence of synthetic and organic fertilizers (EXP1)

Fluxes of N_2_O from EXP 1 are presented on **Fig. 1A**. The main emission peak occurred within the first seven days. Afterward, fluxes from soils amended with OFs returned to background levels, while with ammonium sulfate (AS) N_2_O emissions continued up to Day 14. Consequently, cumulative emissions were calculated for the periods 0–7, 8–14, and 16–48 days, referred to as periods 1, 2 and 3, respectively (**Fig. 1B, blue bars**). During period 1, emissions from animal-based fertilizers (CS and CM) were lower than those from plant-based or mixed fertilizers (O7 and D). In period 2, only AS had significantly higher emissions than all other treatments; in period 3, its emissions matched plant-based but exceeded animal-based fertilizers. The partitioning of N_2_O emissions among the 3 periods was as follows: AS (69/20/11), D (89/0/11), O7 (83/1/17), CM (95/0/5), and CS (100/0/0). In this experiment, the initial ammoniacal N content of each fertilizer was a key determinant driving N_2_O emissions, particularly in treatments with AS, D, and O7 (**Table S2**). Thus, total cumulative emissions were the highest for AS fertilization across all the three periods, resulting in 1992 µg N_2_O-N kg^−1^ of dry soil. Corresponding total cumulative emissions for D, O7, CM and CS treatments were 998, 883, 395 and 127 µg N_2_O-N kg^−1^ of dry soil, respectively, whereas the unfertilized soil (NF) exhibited a low and negative total cumulative emission of −71.4 µg N_2_O-N kg^−1^ of dry soil.

**Fig. 1.**
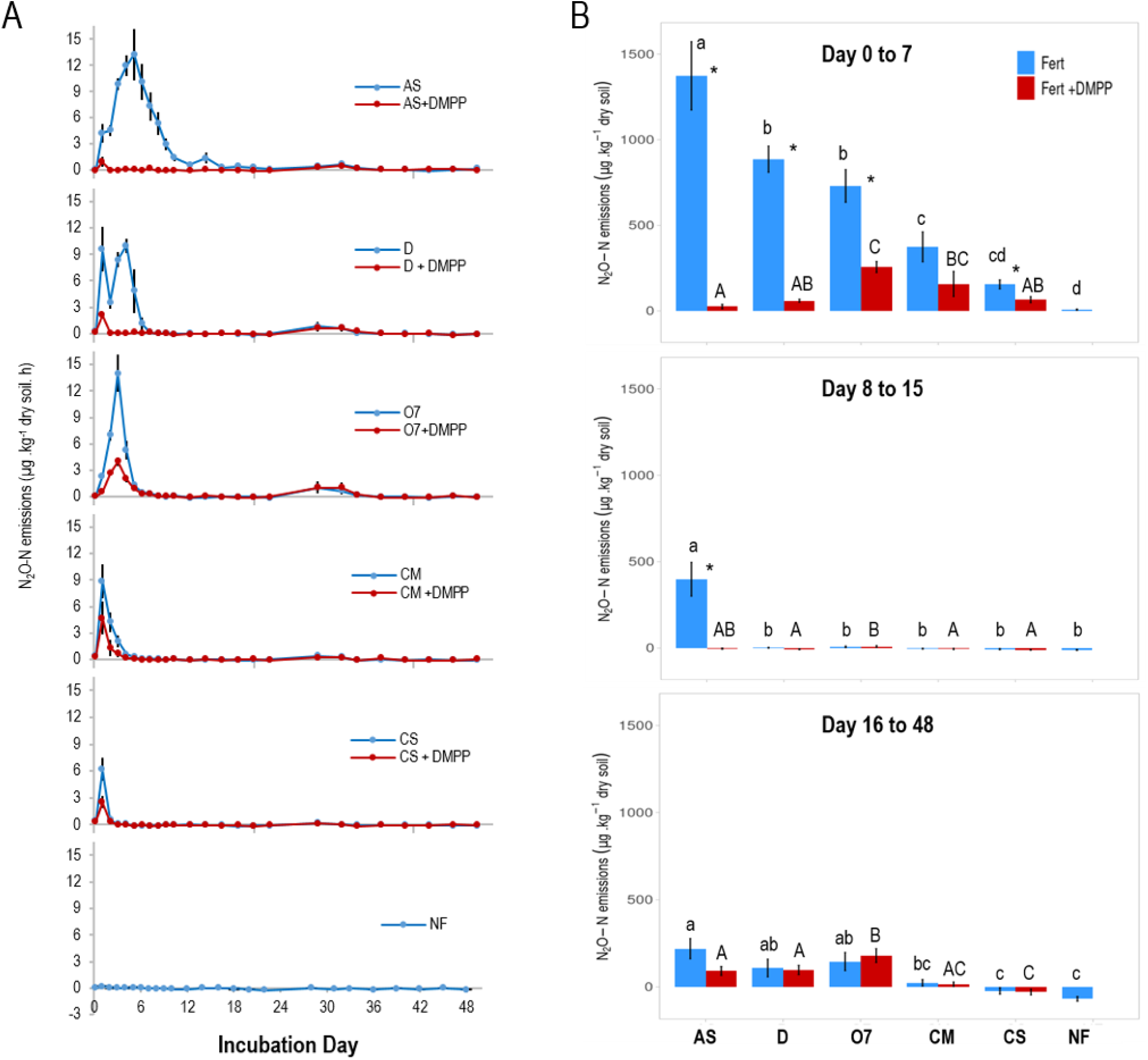
Soil N_2_O fluxes (A) for each fertilization treatment without (blue) and with the addition of DMPP (red). Dotted lines show separation between periods. N_2_O cumulative emissions (B) for the three different periods. Values represent mean ± SE (n = 4). For each period, letters represent significant differences between fertilizer treatments analyzed by Duncan’s test (P<0.05), lower case letters for treatments without the inhibitor and upper-case letters for treatments with DMPP. Asterisk indicates significant differences between same fertilizer with and without DMPP (T-student, p < 0.05).

The addition of DMPP inhibited N_2_O emissions for all fertilizer treatments. DMPP was the most effective when applied with AS, achieving a 94% reduction during period 1, followed by D and O7 treatments, with reductions of 85% and 61%, respectively (**Fig. 1B, red bars**). During period 2, the effect of DMPP was only relevant for AS, as fluxes for all other treatments had returned to background levels. In period 3, DMPP no longer influenced N_2_O emissions and did not significantly reduce the minor delayed N_2_O emission peak observed around Day 31.

### 3.2. Changes in soil mineral N, C and pH (EXP1)

Changes in soil ammonium (NH_4_^+^-N) and nitrate (NO_3_^−^-N) concentrations are shown in **Fig. 2 (blue bars)**. Despite the differing initial NH_4_^+^-N levels among amendments (**Fig. 2A, day 0**), soil NH_4_^+^-N content decreased by Day 7 across all fertilized treatments. For those amended with OFs, NH_4_^+^-N levels even reached values comparable to the non-fertilized treatment by Day 15. Patterns of NO_3_^−^-N dynamics varied among fertilizer type. In treatments with AS and plant-based/mixed organic fertilizers D and O7, NO_3_^−^-N content increased by Day 7 and remained relatively constant until the end of the incubation. On the contrary, in treatments with animal-based fertilizers CM and CS, soil NO_3_^−^-N decreased by Day 7 and remained low up to Day 48. The addition of DMPP (**Fig 2, red bars**) significantly maintained higher NH_4_^+^-N levels in the soil at Day 7 and Day 15 when applied with fertilizers AS, D, and CM. Additionally, DMPP effectively reduced the intermediate nitrification product, nitrite (NO₂⁻^−N^), at Day 7 for AS, D, and O7 (**Fig. S2**). However, DMPP only significantly reduced NO_3_ ^−^-N presence (**Fig. 2**) for AS and CM at Day 7.

**Fig. 2.**
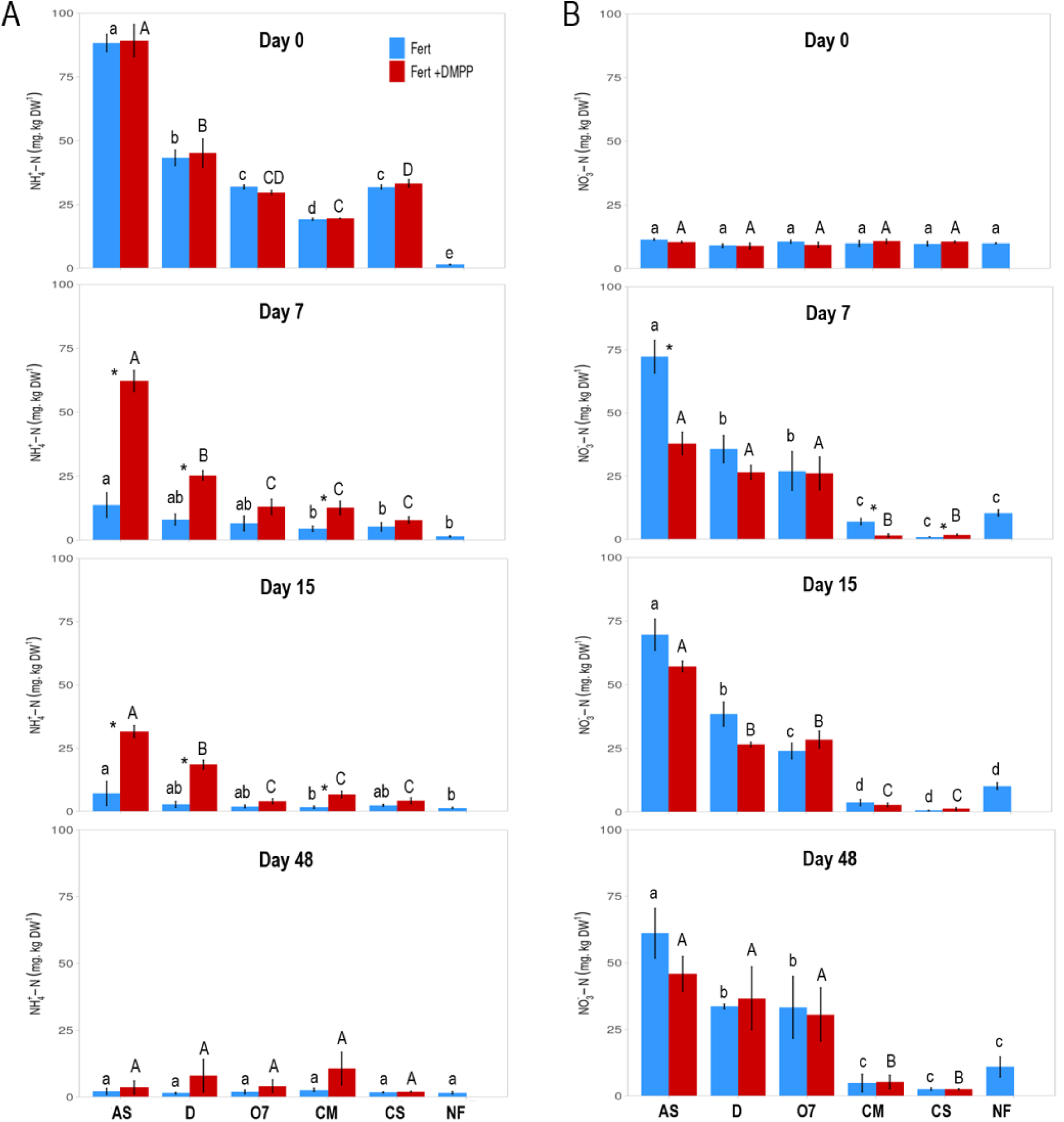
Soil NH_4_^+^ content (A) and NO_3_^−^ content (B) for each fertilizer treatment without (blue) and with the addition of DMPP (red) (EXP1). Values represent mean ± SE (n = 4). For each incubation day, letters represent significant differences between fertilizer treatments analyzed by Duncan’s test (P<0.05), lower case letters for treatments without the inhibitor and upper-case letters for treatments with DMPP. Asterisk indicates significant differences between same fertilizer with and without DMPP (T-student, p < 0.05).

Initial (Day 0) soil dissolved organic carbon (DOC) content was dependent of the fertilization treatment (**Fig. S3**). DOC represents the readily available carbon pool in the soil, which serves as an energy source for microbial activity and influences nutrient cycling (Kalbitz et al., 2000). The highest DOC values were for CS, followed by CM and D; while the addition of AS and O7 did not show any increase in the initial DOC value compared to NF. For animal-based OFs, despite their higher initial C content (**Fig. S1**), DOC contents decreased by Day 7 and remained low until the end of the incubation period. The addition of DMPP did not affect DOC values in any of the treatments. Soil pH responses to chemical reactions within the soil matrix and plays a crucial role in regulating microbial activity. In our study, all fertilizers caused a slight acidification compared to NF (**Fig. S4**). likely due to nitrification process releasing protons. Notably, the presence of DMPP led to higher pH values during the first seven days when applied with AS, possibly by inhibiting ammonia oxidation and delaying proton release. However, when combined with other fertilizers, DMPP had minimal impact on soil pH, suggesting its effect is dependent on the specific composition of the applied N source.

### 3.3. Abundance of N-cycling bacterial and archaeal populations

Total bacterial and archaeal abundances in the incubated soils were not significantly affected by the tested fertilizers compared to NF, except a decrease in archaeal population for CS along the incubation time (**Fig. S5**). Similarly, the initial (Day 0) abundance of marker genes for both nitrifying and denitrifying populations was comparable across treatments (**Fig. 3; Figs. S5–S7)** indicating that subsequent changes in these genes were driven solely by shifts in the indigenous soil microbiota rather than by the introduction of additional microorganisms through organic fertilizers. The abundances of bacterial (AOB) and archaeal (AOA) nitrification genes *amoA* are presented in **Fig. 3**. In absence of DMPP, AOB populations increased in fertilized soils by Day 7 for AS and D treatments and for all fertilized treatments by Day 15, with the highest values observed in soil treated with AS (**Fig. 3A**). At Day 7 and Day 15, DMPP significantly reduced AOB populations to levels comparable to NF for all treatments except CS. At Day 7, the reduction percentages ranged from 72% to 43%, and at Day 15 from 69% to 45%, with the highest efficiencies observed in AS+ and D+ treatments. By Day 48, AOB abundances had returned to initial levels, irrespective of DMPP presence, so the effect of the inhibitor was no longer detectable.

**Fig. 3.**
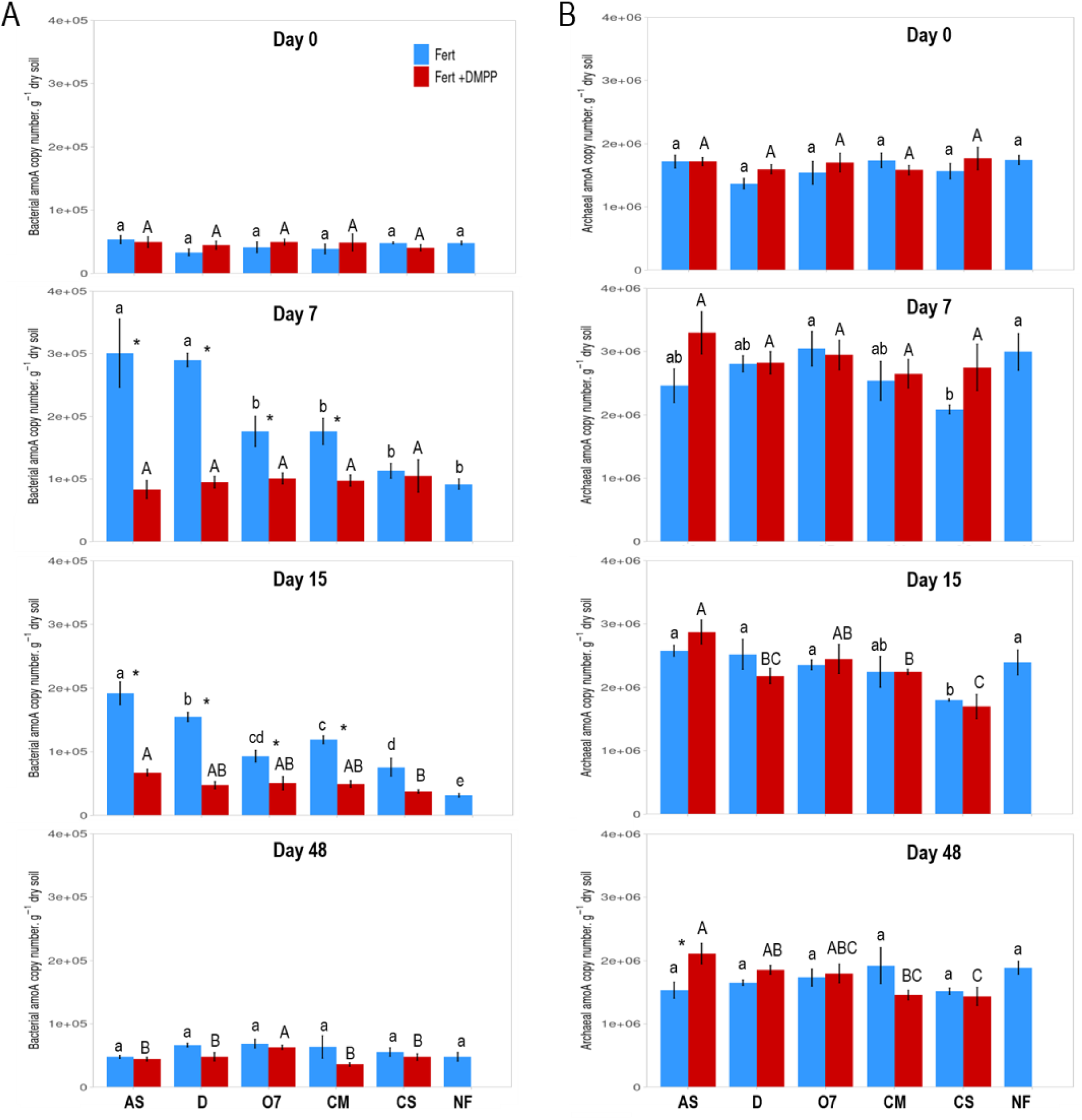
Abundance of ammonia oxidizing bacteria (AOB) (A) and archaea (B) (AOA) determined by *amoA* marker gene at Day 0, 7, 15 and 48 after fertilization (EXP1). Values represent mean ± SE (n = 4). Statistics as in Fig. 2.

AOA population increased after seven days of incubation across all the treatments, including NF (**Fig. 3B**). The reduced AOA abundance in the CS treatment is consistent with the observed decrease in the total archaeal population (**Fig. S5B**). AOA population was more abundant than AOB in the tested soils but was not significantly affected by DMPP application. By Day 48, AOA abundances had returned to background levels observed on Day 0. The denitrifying bacterial population showed minimal changes in response to fertilization compared to NF, and all of them decreased over incubation time (**Figs. S6– S7**). The addition of DMPP did not induce any consistent response in denitrifying bacterial populations. Isolated increases in denitrifying bacterial populations bearing *nirS* and *nirK* were observed, with the most notable increase in *nirK* abundance for the CM treatment at Day 7. In contrast, *nosZI* abundance was significantly reduced for the D treatment at Day 7.

### 3.4. Bacterial community structure and composition under DMPP with mineral and organic inputs

The fertilizer type and time were the main drivers of bacterial community shifts, regardless the presence of DMPP (**Table 1; Tables S7 and S8**). Significant differences between mineral and organic fertilization (**Table S7**) suggest distinct microbial communities develop in response to different nutrient sources. An early divergence in community composition was observed (**Fig. 4A**) following the application of liquid D, and animal-based fertilizers (CM and CS), which later converged by Day 48, suggesting a temporal homogenization of bacterial communities. In contrast, CM and CS remained clearly separated in the constrained ordination throughout the incubation period, indicating that fertilization-driven community structures persisted over time (**Fig. 4B**). Fertilization also strongly shaped the *Nitrosomonadaceae* family (ammonia-oxidizing bacteria, AOB), particularly at early stages (**Table 1**). Indeed, the genus *Nitrospira* was indicative of non-inhibited treatment*s,* pointing to active nitrification up to nitrate formation (**Table S9**). In contrast, this genus was not detected as indicative taxa under the application of DMPP, confirming the effective suppression of the nitrification bacteria.

**Fig. 4.**
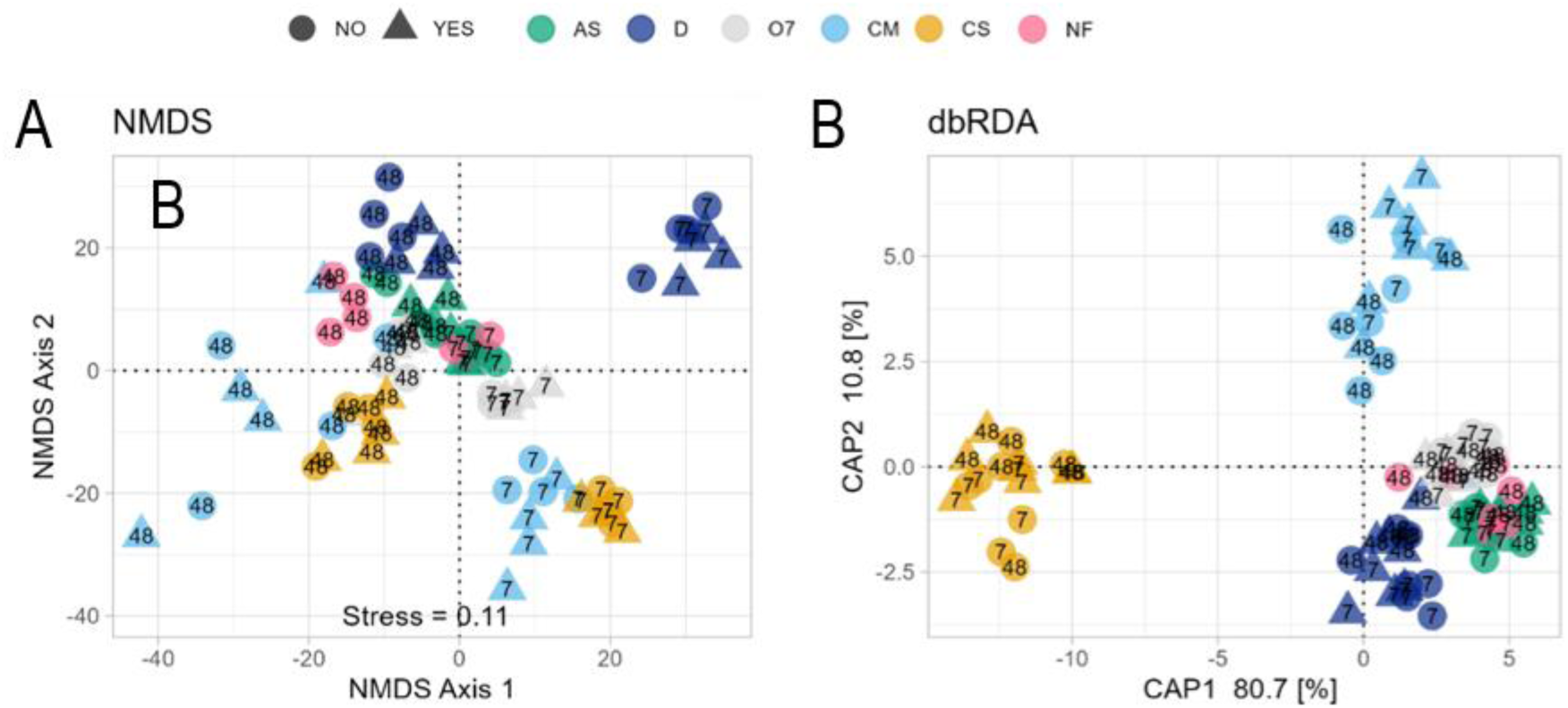
Unconstrained constrained ordination of bacterial community structure as assessed by Non-metric multidimensional scaling (NMDS) and (D) distance-based Redundancy Analysis (dbRDA) with fertilization type as constraining factor. Symbols show DMPP application and colors fertilization type. Labels represent incubation time.

Throughout the incubation period, bacterial community structure showed limited response to DMPP. A marginal impact on overall bacterial structure was only detected at the end of the incubation period under mineral fertilization with AS (**Table 2**). In contrast, DMPP influenced specific nitrifying groups earlier on. Its inhibitory effect on the *Nitrosomonadacea*e family was already evidenced by Day 7 under AS treatment (**Table 2**), indicating early inhibition of ammonia oxidation, consistent with the strong decline in AOB population observed under these conditions (**Fig. 3A**). The rapid N availability following AS application likely stimulated nitrifier activity, making this treatment more susceptible to be affected by DMPP than those amended with organic sources. Additionally, DMPP exerted a general delayed impact over the final step of the nitrification, performed by nitrite-oxidizing bacteria (NOB), part of Nitrospirae class (**Table 1**), possibly reflecting microbial succession and earlier nutrient shifts driven by AOB inhibition.

**Table 2:**
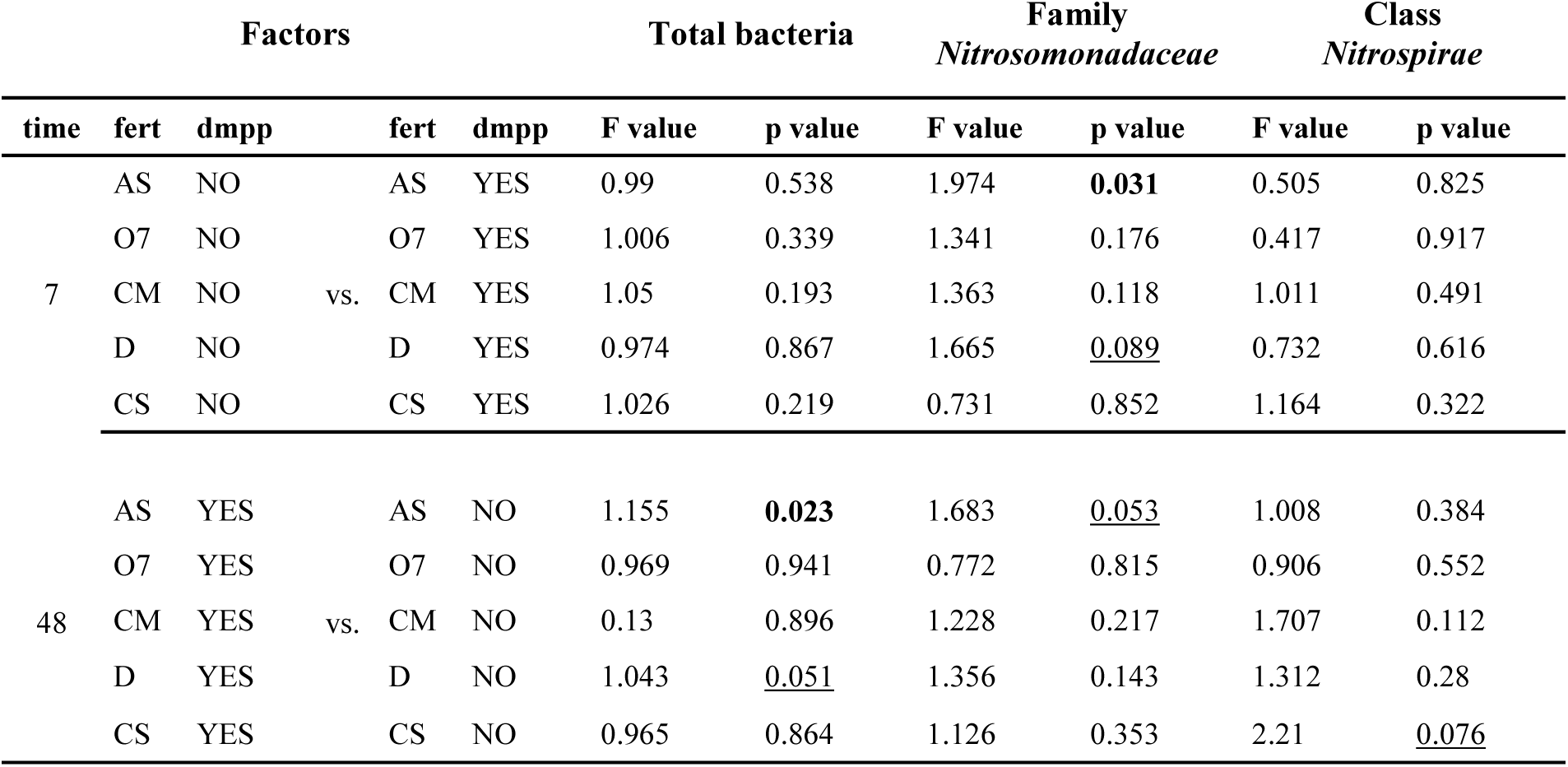
Pairwise comparison of bacterial community structure for total bacteria, the class of *Nitrospira* and the familiy of *Nitrosomonadaceae* as affected by DMPP addition. Bold letters show significant differences at p>0.05, underlined letters show statistical trends P=0.05-0.1

### 3.5. Soil N_2_O emissions from synthetically or biologically inhibited fertilized soil (EXP2)

Cumulative N_2_O emissions from Experiment 2 are presented in **Fig. 5**, and daily fluxes are shown in **Fig. S9.** Background emissions (NC treatment = untreated and non-fertilized control soil) were ca. 0.27 µg N_2_O-N kg^−1^ of dry soil h^−1^, similar to emissions from the SDMPP treatment after AS fertilization (**Fig. S9**). In the absence of soil treatment (NT), the fertilizer resulting in the highest cumulative N_2_O emissions over the 21-days study period was Orgamine 7 (6843 µg N_2_O-N kg^−1^ of dry soil), followed by Digestate (D) (3225 µg N_2_O-N kg^−1^ of dry soil), and AS (1401 µg N_2_O-N kg^−1^ of dry soil). Regardless of the fertilizer type, N_2_O emissions were significantly reduced by DMPP application (SDMPP), with reductions of 88% for AS, 92% for D, and 80% for O7.

**Fig. 5.**
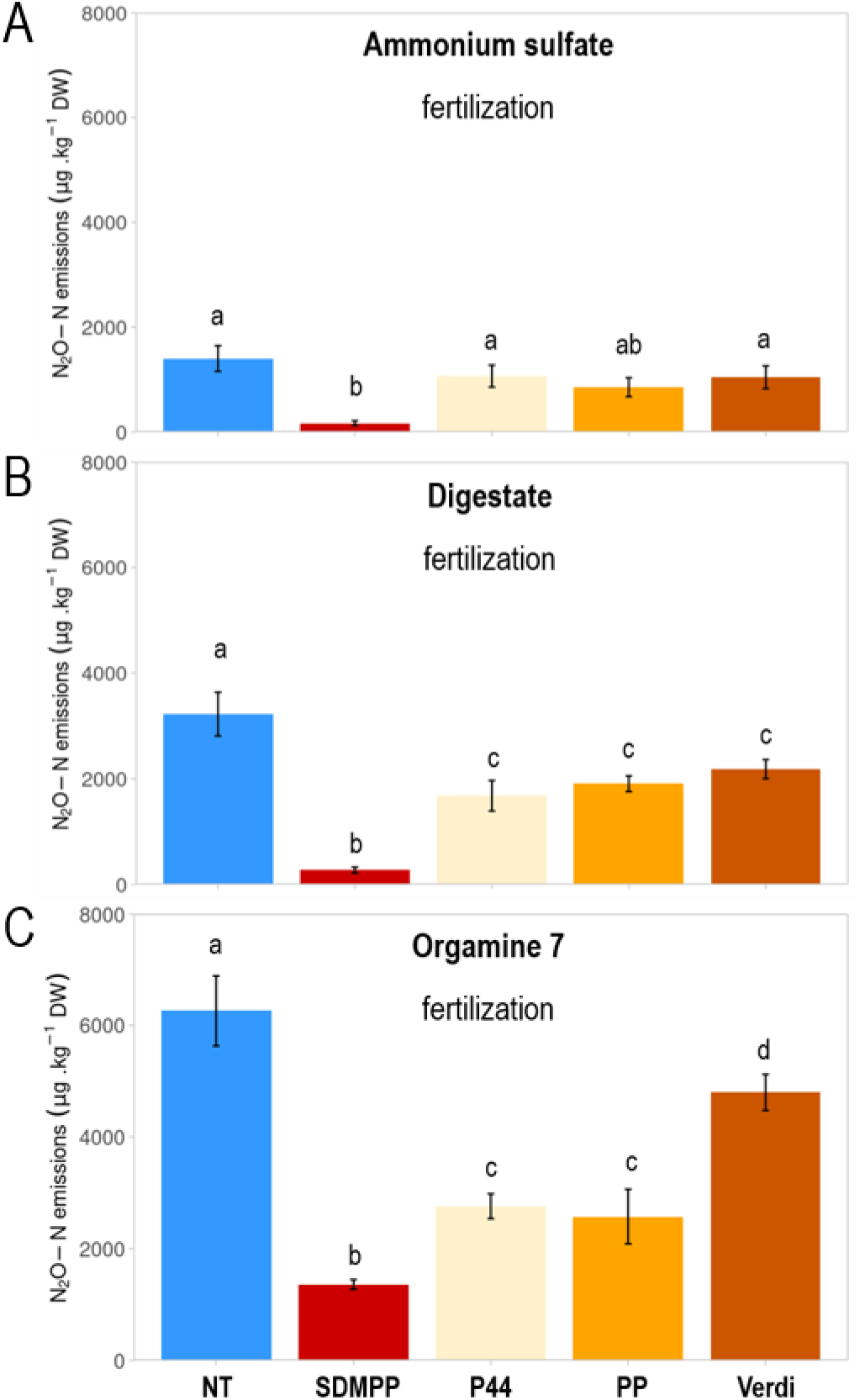
N_2_O cumulative emissions for 21-days incubation period of each soil treatment (EXP2). Incubations consisted in not treated soil (NT), soil treated with DMPP (SDMPP), and soil collected from the rhizosphere of wheat cv. Persia 44 (P44), white mustard cv. Pole Position (PP) and cv. Verdi (VERDI). Soils were incubated with fertilizers (A) ammonium sulfate (AS), (B) liquid digestate (D) and (C) Orgamine 7 (O7). For each fertilizer, letters represent significant differences between soil treatments analyzed by Duncan’s test (P<0.05).

Following application of mineral fertilizer AS (**Fig. 5A**), none of the rhizospheric soils of wheat (Persia 44 landrace, P44) nor of white mustard (cv. Pole Position, PP or cv Verdi) could significantly reduce N_2_O emission as compared to the non-treatment soil control (NT). In contrast, the ability of those plants to produce BNI compounds became only evident with organic fertilizers (OFs) (**Fig. 5B-C),** where N_2_O emissions were significantly reduced compared to NT soils. Under D fertilization, all three BNI-producing plants exhibited a similar reduction of about 40 % in N_2_O emission levels (**Fig. 5B**). With O7, rhizosphere soils from P44 and PP plants showed even stronger inhibition, reducing N_2_O emissions by 60% and 62%, respectively (**Fig. 5C**).

### 3.6. Effect of BNI presence in the soil over mineral N contents (EXP2)

NH_4_ ^+^-N and NO_3_ ^−^-N concentrations in soil collected from the plant cultivation phase (from pots with or without plants) are presented in **Fig. 6A and E**. After four weeks of irrigation with ammonium sulfate (AS) solution, NH_4_ ^+^-N levels in NT and plant-treated pots ranged from 16 to 17.2 mg kg^−1^ of dry soil, whereas in soil irrigated with AS+DMPP, NH_4_ ^+^-N concentration remained at 25.7 mg kg^−1^ dry soil by the end of the treatment period. By contrast, NO_3_ ^−^-N concentration was the highest in NT pots, decreasing from 33.7 to 10.7 kg^−1^ dry soil when AS had been supplied together with DMPP. The growth of wheat and white mustard plants reduced rhizospheric soil NO_3_ ^−^-N levels to background values c.a. 1.7 kg^−1^ dry soil.

**Fig. 6.**
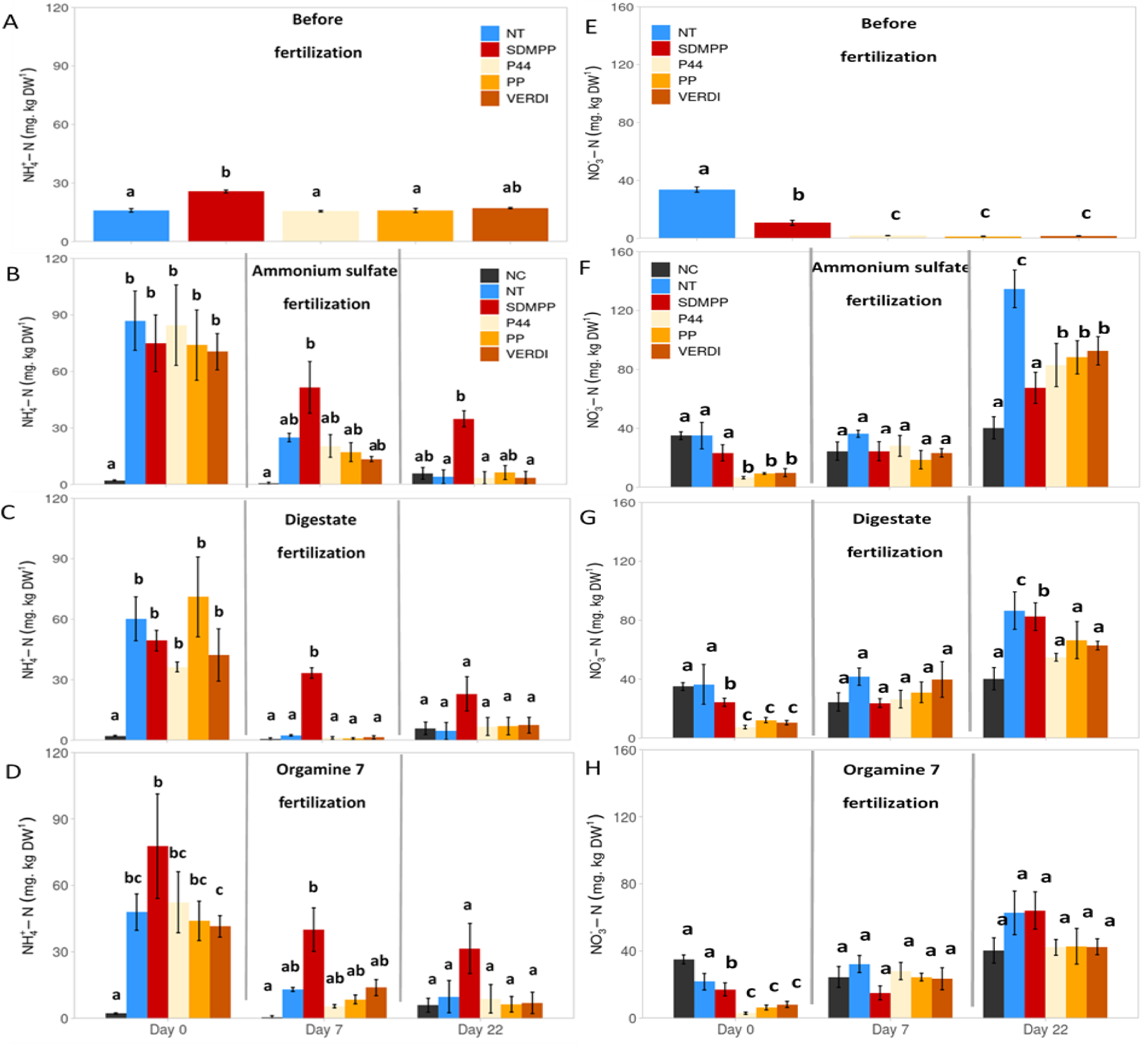
Soil mineral N content (NH_4_^+^ and NO_3_^−^) content in incubation pots (EXP 2) with not treated soil (NT), soil treated with DMPP (SDMPP), and rhizospheric soil from wheat (P44) and white mustard (VERDI and PP). Mineral N was determined before (upper panels A and E) and after the addition of the fertilizer fertilizers (B,F) ammonium sulfate (AS), (C,G) liquid digestate (D) and (D,H) Orgamine 7 (O7). NC, untreated and non-fertilized control soil. Values represent mean ± SE (n = 4). For each incubation day and fertilizer, letters represent significant differences between soil treatments analyzed by Duncan’s test (P<0.05).

After setting up the incubation pots, NH_4_^+^-N and NO_3_^−^-N concentrations were measured immediately after applying mineral (AS, **Figs 6 B and F**) or organic (D or O7, **Figs C-D** and **G-H**) fertilizers (Day 0), and later on Day 7 and Day 21. Initial NH_4_^+^-N values at Day 0 directly reflected the NH_4_^+^ content (%) of each fertilizer, with the mineral N source (AS) showing the highest concentration (**Table S2**). By Day 7, NH_4_^+^-N concentration decreased sharply for NT and rhizospheric soils, approaching those of the non-fertilized control (NC). Meanwhile, the synthetic inhibitor treatment (SDMPP) efficiently preserved high NH_4_^+^-N concentrations in both mineral and OFs (**Fig. 6B-D**), consistent with observations from EXP 1. Regarding NO_3_^−^, its concentration remained unchanged at Day 7 in untreated soils (**Fig. 6F-G**), but it increased significantly compared to NC at Day 21, particularly with AS and D fertilization. In SDMPP treatment, this increase was observed when incubated with AS, where NO_3_^−^-N levels remained at half those observed in NT soil. In rhizospheric soil from BNI-producing plants, NO_3_^−^ content also increased by Day 21 but remained lower than in NT soil especially with mineral fertilizer, mirroring the effect of DMPP. For rhizospheric soils incubated with OFs, NO_3_^−^-N concentrations were generally lower than in NT soil, with a significant reduction observed in P44 rhizospheric soil fertilized with digestate (D).

## 4. Discussion

### 4.1. Fertilizer type influences N cycling and microbial community dynamics

The transition toward sustainable farming practices is essential for improving environmental health, enhancing soil quality, and supporting biodiversity. Extending agricultural land use under organic farming is one of the cornerstones of the European Green Deal’s agricultural objectives, aiming for climate neutrality by 2030 (Guyomard, Bureau et al., 2020). Our study demonstrated that fertilizer type significantly affected N_2_O emissions, N mineral status and nitrifier communities. Mineral fertilizers are generally linked to higher emissions (Aguilera et al., 2013; Sun et al., 2024; Wu et al., 2024), though the contrary has been reported in other studies (Petersen et al., 2023; Li et al., 2024). Indeed, in Experiment 1, mineral fertilizer (AS) generated the highest N_2_O emissions, followed by plant-based liquid digestate (D) or the Orgamine7 (O7) pellets. These fertilizers produced higher total accumulated emissions than animal-based fertilizers (CM and CS). In contrast, in Experiment 2, OFs showed higher N_2_O emission rates than the mineral N source, with Orgamine7 showing the highest values, suggesting that the results could have been influenced by the soil properties (that was mixed with sand for EXP2) or the specific properties of the fertilizer batch used in each incubation experiment. (Chen et al., 2013; Cayuela et al., 2017; Li et al., 2023). In any case, the increase in soil NO_3_ ^−^-N concentration with time observed in soils fertilized with AS, D or O7 in both experiments (**Fig. 2B**, **Fig. 6F-H**) implies that nitrification was likely the main process contributing to the high N_2_O emissions peak. This is further supported by the increased abundance of AOB by Day 7 (**Fig. 3A**), which aligned with the N_2_O emission (**Fig. 1**), particularly after the addition of AS and D. Interestingly, although AOB populations were still abundant across all fertilized treatments at Day 15, only mineral fertilizer (AS) exhibited sustained N_2_O emissions beyond Day 7 (**Fig. 1A**), supported by the high NH_4_ ^+^ availability **(Fig. 2A).**

By the end of the incubation, the overall bacterial population declined to pre-fertilization values across all the treatments (**Figs. S5**), including nitrifying and denitrifying bacteria (**Fig. 3, Fig. S6-7)**, likely due to the depletion of soil nutrients, including NH_4_^+^ substrate (**Fig. 2A**). Interestingly, the composition of the soil bacterial communities varied significantly over time (**Table 1** and **S7**). This pattern aligns with previous findings showing that sustained AS fertilization can lead to changes in bacterial community composition in grassland (Corrochano-Monsalve et al., 2021) and arable soils (Zhou et al., 2015; Lori et al., 2023). Furthermore, the bacterial community composition differed significantly between mineral fertilization and the rest of the treatments, regardless the incubation time (**Table S8**), suggesting that application of a high NH_4_^+^ source exerts a strong selective pressure on microbial communities, probably favoring taxa adapted to high NH_4_^+^ availability. Unlike bacteria, total archaeal population exhibited limited changes during incubation (**Fig. S5B**), and AOA were less responsive to fertilization or fertilizer type compared to AOB (**Fig. 3B**). These findings are consistent with studies showing that AOB are the primary contributors to nitrification-driven N_2_O emissions in alkaline soils (Lei et al., 2022) and N-rich environments (Carey et al., 2016; Ouyang et al., 2017).

The magnitude and duration of N_2_O emissions from manure depend on its composition and quality, regarding not only their N content but also the C/N ratio, pH or water content (Valkama et al., 2024). Thus, in our study, OFs with the highest C content and C/N ratios, namely CS and CM (Table S2), produced the lowest N_2_O emissions (**Fig. 1**), as the increased C availability may promote microbial N immobilization (Mooshammer et al., 2014). These results are in line with previous findings where fertilizers with lower C/N ratios increased N availability and stimulate N_2_O production associated to nitrification (Chen et al., 2013). Besides, the simultaneous consumption of not only NH_4_ ^+^, but also NO_3_ ^−^ in soil incubated with animal-manure (**Fig. 2B)** would suggest the implication of denitrification. However, in our study, denitrifier populations were unaffected by fertilizer addition (**Figs. S6-S7**), indicating that the aerated conditions (60% WFPS) and low available C after Day 7 (**Fig. S3**) did not favor their growth, which relies on anaerobic conditions and sufficient C (Butterbach-Bahl et al., 2013).

### 4.2. DMPP efficiently stabilizes N_2_O losses from organic fertilizers

The effectiveness of DMPP a depends on soil conditions and fertilizer type (Yang et al., 2016; Lei et al., 2022). In our study, DMPP highly reduced N_2_O emissions by 49%–94%, in line with the meta-analysis by Lei et al. (2022) and surpassing the 12%–16% mean reductions reported in the meta-analysis by Tufail et al. (2023) for organic and chemical fertilization respectively. This higher efficiency can be attributed to the optimized incubation conditions (60% WFPS) that favor nitrification and enhance DMPP’s impact. This was also demonstrated by the meta-analysis conducted by Soares et al. (2023), with effective conditions ranging from 50 to 75 % WFPS, in which nitrification is the predominant process contributing to N_2_O emissions, (Tufail et al., 2023). The observed impact of DMPP on AOB population (**Fig. 3A**) further supports its efficiency in the present study, and is consistent with the reported specificity of DMPP for bacterial nitrifiers in alkaline soils (Kleineidam et al., 2011; Chen et al., 2015; Torralbo et al., 2017, 2017; Fan et al., 2019; Zhou et al., 2020).

DMPP was most effective in reducing nitrification in AS and D treatments, amendments with the highestN_2_O emissions rates, for which the inhibitor was able to maintain increased soil NH_4_^+^ levels up to Day 15 (**Fig. 2A**). The increased availability of this easily assimilable N source is crucial for efficient plant growth (Coskun et al., 2017; Subbarao and Searchinger, 2021). Thereby, our results suggest that liquid digestate (D) could be a very suitable organic amendment to design optimized fertilization strategies aimed at enhancing plant NUE while mitigating gaseous N losses. However, the effectiveness of DMPP over soil nitrification diminished over time (**Fig. 1**, **Fig. 3A**), as NH_4_^+^ and NO_3_^−^ levels in DMPP-treated soils converged with those of uninhibited soils by the end of the incubation (**Fig. 2**). The direct effect of DMPP over soil nitrifying bacteria (AOB) usually ranges between two and six weeks either in microcosms studies or when applied in the field (Yang et al., 2016b; Tufail et al., 2023b; Yin et al., 2023), which constitutes one of the main limitations of using synthetic NIs (Coskun et al., 2017; Sadhukhan et al., 2022).

In our incubation study, DMPP was less effective with animal-based fertilizers. The literature reports contrasting results regarding the effectiveness of DMPP in mitigating N₂O emissions, with its performance being either superior or inferior when applied with organic manures compared to mineral fertilizers (Aguilera et al., 2013; Petersen et al., 2023; Li et al., 2024; Sun et al., 2024; Wu et al., 2024). It can be affected by the potentially increased NH_3_ losses in presence of DMPP (Wu et al., 2021). In our case, the depletion of NO_3_ ^−^ levels in CM and CS treatments, regardless the presence of the inhibitor (Fig. 2B), as well as the high DOC consumption (**Supplementary Fig. S3**), common indicator of C utilization from microbial activity, suggested that DMPP may have promoted denitrification over nitrification. Long-term studies (>40 days) have shown that while DMPP effectively suppresses nitrification initially, it may also influence denitrifer communities over time, potentially increasing *nosZ* gene abundance indicative of complete denitrification, reducing thus the N_2_O levels (Barrena et al., 2017; Torralbo et al., 2017; Chen et al., 2019; Huérfano et al., 2022). Nevertheless, despite the initial steps of denitrification (by means of *nirS* and *nirK* gene abundances) were sporadically affected by DMPP, *nosZ I* levels did not positively respond to the presence of the inhibitor (**Supplementary Figs. S6-7**), further indicating that the effect of DMPP was limited to nitrification in this study.

### 4.3. DMPP targets nitrifying community without affecting total soil bacterial community

The bacterial community structure remained largely unaffected by DMPP throughout the incubation period (Fig. 4, Table 1), aligning with previous studies where DMPP was applied to alkaline soils fertilized with urea (Zhang et al., 2017; Bachtsevani et al., 2021; Wang et al., 2022) or to acidic grasslands fertilized with AS (Corrochano-Monsalve et al., 2021). Similarly, DCD did not alter bacterial community structure when applied with swine slurry (Suleiman et al., 2016). In contrast, the nitrification inhibitor DMPSA tended to reduce both diversity and richness, although its effect was dependent on soil water content (Corrochano-Monsalve et al., 2020, 2021). The persistence of temporal changes in β-diversity across fertilizer treatments (Fig. 4), despite DMPP application, further highlights that the inhibitor did not influence overall bacterial diversity in this study.

Notably, DMPP specifically targeted members of the *Nitrosomonadaceae* family in the early stages of the incubation (Table 1 and 2), and its effects extended to *Nitrospira* populations toward the end of the experimental period. While uninhibited fertilized conditions were associated with the *Nitrospira* genus (Table S9), which responds to elevated soil NH_4_^+^ levels (Yuan et al., 2023) and have been shown to be abundant also in organic arable systems (Zhang et al., 2019), samples treated with DMPP showed no correlation with this genus. This further confirms the effectiveness of the inhibitor to suppress the nitrification pathway. A comparable reduction in Nitrospirae phylum by DMPP was reported by Corrochano-Monsalve et al. (2021) in soils incubated with AS, linking the inhibition not only to the reduced nitrite oxidation but also to the suppression of ammonia and hydroxylamine oxidation (Daims et al., 2015). In contrast, other taxa, including those linked to denitrification, remained unaffected by fertilizer type or DMPP, mirroring the stability observed in marker gene abundances for denitrifying bacteria (**Supplemenary Figs S6 and S7**). This selective inhibition highlights the specificity of DMPP’s action on nitrifiers under our established incubation conditions, without broadly altering other microbial groups. However, DMPP’s effect is transient, and once its inhibitory action fades, the accumulated NH₄⁺ can be rapidly oxidized, potentially triggering enhanced nitrification and subsequent denitrification as evidenced by the already equal NO₃⁻ concentrations across all treatments by Day 15 (Fig. 2). This delayed N transformation represents a potential environmental trade-off and has been linked to increased post-harvest N₂O emissions, as observed by Scheer et al. (2017) in vegetable systems. These findings emphasize the need to consider the timing of DMPP degradation and residual N availability, as mismanagement could offset its initial mitigation benefits and contribute to unintended environmental impacts.

### 4.4. BNI is a potential alternative to DMPP

Since the effectiveness of DMPP is generally time-limited, complementary approaches are essential to achieve sustainable and long-term control over the soil nitrification. The research on the biological inhibition of soil N_2_O emissions has been predominantly associated with a limited number of high-value crops (sorghum, rice, and maize) and mineral nitrogen sources or urea (reviewed in Wang et al., 2021; Saud et al., 2022; Zhang et al., 2022). To expand this capacity, new strategies are needed. One approach is the exploration of BNI traits in wild relatives of staple crops. For example, *leymus racemosu,* a perennial grass species related to wheat that can inhibit soil nitrification, has had its BNI-associated genetic material successfully transferred to elite wheat varieties (Subbarao et al., 2021; Bozal-Leorri et al., 2022). Alternatively, Persian wheat landraces identified by O’Sullivan et al. (2016), such as the cultivar Persia 44, have shown nitrification inhibition (Jáuregui et al., 2023), although its potential to reduce soil N_2_O emissions remains untested. Another strategy is to incorporate cover crops with BNI capacity into agronomic rotations to reduce soil nitrification for the subsequent target crops. Sorghum is the most studied crop in this context, with evidence that its inclusion in rotations can reduce nitrogen losses (Bozal-Leorri et al., 2021, 2023; Ma et al., 2023; Zhang et al., 2023; Vega-Mas et al., 2024). However, the warm temperature requirements of sorghum limit its suitability in cooler climates. In northern regions, identifying herbaceous species with BNI ability that are better adapted to cooler climates, such as certain *Brassicaceae* species like mustard and rapeseed, could provide viable alternatives, as their incorporation into soil has been shown to reduce nitrification activity and promote NH_4_^+^ accumulation (Bending and Lincoln, 2000; Brown and Morra, 2009; Heuermann et al., 2019).

Our incubation experiment (EXP 2) proved the potential of rhizospheric soil from wheat landrace Persia 44 and white mustard (PP and Verdi) cultures to reduce soil N_2_O emissions for at least nine days following organic fertilization (**Fig. S9**). Interestingly, our findings from EXP 2 highlighted that, similarly to synthetic NIs, the efficiency of rhizospheric soils in reducing N_2_O emissions appears to depend on the type of fertilizer applied (**Fig. 5**). Notably, rhizospheric soils demonstrated greater effectiveness in mitigating N_2_O emissions when fertilized with OFs, whether it was liquid D or O7 pellets. This is particularly remarkable for the application of O7, as the reduction factors of 50 % and 55 % observed in the rhizospheric soils of white mustard (Pole Position, PP) and wheat landrace Persia 44 (P44), respectively, were nearly comparable to those achieved with DMPP. In the case of rhizospheric soil incubated with D, the reduced nitrifying conditions of the soil did not only limit N_2_O emissions (Fig. 5) but also maintain reduced NO_3_ ^−^ levels in the soil **(Fig. 6B**), that could potentially diminish leaching losses in an agronomical context. The combination of biological mitigation tools together with revalorized organic fertilizers provides a step forward in the search for an all-organic agriculture that is both efficient and environmentally friendly.

## 5. Conclusion and outlook

Our study emphasizes the importance of considering fertilizer types and composition in shaping N cycling and implementing mitigation strategies to control N losses form agroecosystems. Fertilizers with higher ammoniacal N content, such as mineral AS and plant-based liquid digestate (D), promoted nitrification more strongly, resulting in higher total N_2_O emissions compared to animal-based fertilizers. DMPP proved effective in reducing early nitrification and associated N₂O peaks, particularly under mineral and plant-based fertilization, while maintaining high soil NH_4_^+^ levels for at least two weeks. In contrast, the higher C/N ratios of the animal-sourced OFs contributed to lower baseline N_2_O emissions, making them less efficiently stabilized with DMPP. Although DMPP consistently inhibited bacterial nitrification from all N amendments, its effect specifically targeted nitrifyng bacteria, without impacting denitrifiers or the broader soil bacterial community. However, this inhibition was transient, diminishing after two weeks, underscoring a key limitation of synthetic inhibitors. To address these temporal constraints, our findings support the potential of biologically driven mitigation. Rhizospheric soils enriched by BNI-active plant species, such as wheat landrace (Persia 44) and white mustard (cv. PP and cv. Verdi), successfully suppressed nitrification and associated N_2_O losses under organic fertilization, in some cases matching the performance of DMPP. This suggests that BNI could serve as a robust, longer-term complement or alternative to synthetic inhibition, especially in systems aiming for reduced external inputs. Further refinement of the integration of plant-based inhibition with revalorized OFs, such as liquid D or organic pellets (O7), represents a promising step toward fully organic farming systems that balance agronomic efficiency with environmental sustainability.

## 6. Acknowledgements

The authors would like to acknowledge the financial support of the Walloon Region of Belgium (Wallonie agriculture SPW) grant agreement GAIN (Project D31-1378-/S1) and the SusCrop-ERA-NET Program (Project: Catch-BNI). I.V.M. acknowledges the Basque Government for her postdoctoral fellowship (POS-2018-1-005). H.M.K and C.T acknowledge the SNF and FNRS for joint funding of the PlantaGO project (grant number. 310030L_220103/1). The following organisation are also acknowledged for providing organic fertilizers: S.C. Biogaz Du Haut Geer (Gaëtan de Seny) for the liquid digestate (D), CRA-W (Dr Bernard Godden) for the two animal-based fertilizers (CS and CM) and Fayt-Carlier for Orgamine 7 (O7). The company DSV (Deutsche Saatveredelung) is acknowledged for providing the seeds of both white mustard varieties. Australian Grains Genebank (AGG) is acknowledged for providing the seeds of wheat. We acknowledge the help of Jean-Claude Walser in performing bioinformatic analysis at the Euler Scientific Compute Cluster at ETH Zürich.

## 7. Appendix A. Supplementary data

## References

Aguilera, E., Lassaletta, L., Sanz-Cobena, A., Garnier, J., Vallejo, A., 2013. The potential of organic fertilizers and water management to reduce N2O emissions in Mediterranean climate cropping systems. A review. Agriculture, Ecosystems & Environment 164, 32–52. 10.1016/j.agee.2012.09.006

Bachtsevani, E., Papazlatani, C. V, Rousidou, · Constantina, Lampronikou, E., Menkissoglu-Spiroudi, U., Nicol, G.W., Karpouzas, D.G., Papadopoulou, E.S., 2021. Effects of the nitrification inhibitor 3,4-dimethylpyrazole phosphate (DMPP) on the activity and diversity of the soil microbial community under contrasting soil pH. Biology and Fertility of Soils 1, 3. doi:10.1007/s00374-021-01602-z

Barrena, I., Menéndez, S., Correa-Galeote, D., Vega-Mas, I., Bedmar, E.J., González-Murua, C., Estavillo, J.M., 2017. Soil water content modulates the effect of the nitrification inhibitor 3,4-dimethylpyrazole phosphate (DMPP) on nitrifying and denitrifying bacteria. Geoderma 303, 1–8. doi:10.1016/j.geoderma.2017.04.022

Benckiser, G., christ, E., Herbert, T., Weiske, A., Blome, J., Hardt, M., 2013. The nitrification inhibitor 3,4-dimethylpyrazole-phosphat (DMPP) - quantification and effects on soil metabolism. Plant and Soil 371. doi:10.1007/sl1104-1664-6

Bending, G.D., Lincoln, S.D., 2000. Inhibition of soil nitrifying bacteria communities and their activities by glucosinolate hydrolysis products. Soil Biology and Biochemistry 32, 1261–1269. doi:10.1016/S0038-0717(00)00043-2

Bozal-Leorri, A., Corrochano-Monsalve, M., Arregui, L.M., Aparicio-Tejo, P.M., González-Murua, C., 2021. Biological and synthetic approaches to inhibiting nitrification in non-tilled Mediterranean soils. Chemical and Biological Technologies in Agriculture 8, 1–12. doi:10.1186/S40538-021-00250-7/FIGURES/4

Bozal-Leorri, A., Corrochano-Monsalve, M., Arregui, L.M., Aparicio-Tejo, P.M., González-Murua, C., 2023. Evaluation of a crop rotation with biological inhibition potential to avoid N2O emissions in comparison with synthetic nitrification inhibition. Journal of Environmental Sciences (China) 127, 222–233. doi:10.1016/j.jes.2022.04.035

Bozal-Leorri, A., Subbarao, G. V., Kishii, M., Urmeneta, L., Kommerell, V., Karwat, H., Braun, H.J., Aparicio-Tejo, P.M., Ortiz-Monasterio, I., González-Murua, C., González-Moro, M.B., 2022. Biological nitrification inhibitor-trait enhances nitrogen uptake by suppressing nitrifier activity and improves ammonium assimilation in two elite wheat varieties. Frontiers in Plant Science 13. doi:10.3389/FPLS.2022.1034219

Brown, P.D., Morra, M.J., 2009. Brassicaceae Tissues as Inhibitors of Nitrification in Soil. Journal of Agricultural and Food Chemistry 57, 7706–7711. doi:10.1021/jf901516h

Butterbach-Bahl, K., Baggs, E.M., Dannenmann, M., Kiese, R., Zechmeister-Boltenstern, S., 2013. Nitrous oxide emissions from soils: how well do we understand the processes and their controls? Philosophical Transactions of the Royal Society of London. Series B, Biological Sciences 368, 20130122. doi:10.1098/rstb.2013.0122

Cao, Q., Sun, X., Rajesh, K., Chalasani, N., Gelow, K., Katz, B., Shah, V.H., Sanyal, A.J., Smirnova, E., 2021. Effects of Rare Microbiome Taxa Filtering on Statistical Analysis. Frontiers in Microbiology 12, 11:607325. doi:10.3389/fmicb.2020.607325

Carey, C.J., Dove, N.C., Beman, J.M., Hart, S.C., Aronson, E.L., 2016. Meta-analysis reveals ammonia-oxidizing bacteria respond more strongly to nitrogen addition than ammonia-oxidizing archaea. Soil Biology and Biochemistry 99, 158–166. doi:10.1016/j.soilbio.2016.05.014

Cassman, K.G., Dobermann, A.R., Walters, D.T., 2002. Agroecosystems, Nitrogen-use Efficiency, and Nitrogen Agroecosystems, Nitrogen-use Efficiency, and Nitrogen Management Management.

Cayuela, M.L., Aguilera, E., Sanz-Cobena, A., Adams, D.C., Abalos, D., Barton, L., Ryals, R., Silver, W.L., Alfaro, M.A., Pappa, V.A., Smith, P., Garnier, J., Billen, G., Bouwman, L., Bondeau, A., Lassaletta, L., 2017. Direct nitrous oxide emissions in Mediterranean climate cropping systems: Emission factors based on a meta-analysis of available measurement data. Agriculture, Ecosystems & Environment 238, 25–35. doi:10.1016/J.AGEE.2016.10.006

Chadwick, D.R., John, F., Pain, B.F., Chambers, B.J., Williams, J.R., 2000. Plant uptake of nitrogen from the organic nitrogen fraction of animal manures: a laboratory experiment. The Journal of Agricultural Science 134, 159–168.

Chaves, B., Opoku, A., De Neve, S., Boeckx, P., Cleemput, O., Hofman, G., 2006. Influence of DCD and DMPP on soil N dynamics after incorporation of vegetable crop residues. Biology and Fertility of Soils 43, 62–68. doi:10.1007/s00374-005-0061-6

Chen, H., Li, X., Hu, F., Shi, W., 2013. Soil nitrous oxide emissions following crop residue addition: a meta-analysis. Global Change Biology 19, 2956–2964. doi:10.1111/GCB.12274

Chen, H., Yin, C., Fan, X., Ye, M., Peng, H., Li, T., Zhao, Y., Wakelin, S.A., Chu, G., Liang, Y., 2019. Reduction of N2O emission by biochar and/or 3,4-dimethylpyrazole phosphate (DMPP) is closely linked to soil ammonia oxidizing bacteria and nosZI-N2O reducer populations. Science of the Total Environment 694, 133658. doi:10.1016/j.scitotenv.2019.133658

Chen, Q., Qi, L., Bi, Q., Dai, P., Sun, D., Sun, C., Liu, W., Lu, L., Ni, W., Lin, X., 2015. Comparative effects of 3,4-dimethylpyrazole phosphate (DMPP) and dicyandiamide (DCD) on ammonia-oxidizing bacteria and archaea in a vegetable soil. Applied Microbiology and Biotechnology 99, 477–487. doi:10.1007/s00253-014-6026-7

Chynoweth, D.P.; Owens, J.M.; Legrand, R., 2001. Renewable Methane from Anaerobic Digestion of Biomass. Renew. Energy 22, 1–8. doi: 10.1016/S0960-1481(00)00019-7.

Corrochano-Monsalve, M., González-Murua, C., Estavillo, J.M., Estonba, A., Zarraonaindia, I., 2020. Unraveling DMPSA nitrification inhibitor impact on soil bacterial consortia under different tillage systems. Agriculture, Ecosystems and Environment 301, 107029. doi:10.1016/j.agee.2020.107029

Corrochano-Monsalve, M., González-Murua, C., Estavillo, J.M., Estonba, A., Zarraonaindia, I., 2021. Impact of dimethylpyrazole-based nitrification inhibitors on soil-borne bacteria. Science of The Total Environment 792, 148374. doi:10.1016/J.SCITOTENV.2021.148374

Coskun, D., Britto, D.T., Shi, W., Kronzucker, H.J., 2017. How Plant Root Exudates Shape the Nitrogen Cycle. Trends in Plant Science 22, 661–673. doi:10.1016/j.tplants.2017.05.004

Daims, H., Lebedeva, E. V., Pjevac, P., Han, P., Herbold, C., Albertsen, M., Jehmlich, N., Palatinszky, M., Vierheilig, J., Bulaev, A., Kirkegaard, R.H., Von Bergen, M., Rattei, T., Bendinger, B., Nielsen, P.H., Wagner, M., 2015. Complete nitrification by Nitrospira bacteria. Nature 528, 504. doi:10.1038/NATURE16461

De Cáceres, M., Legendre, P., Wiser, S. K., Brotons, L. 2012. Using species combinations in indicator value analyses. Methods in Ecology and Evolution 3, 973–982. doi:10.1111/j.2041-210X.2012.00246.x

Edgar, R.C., 2010 Search and clustering orders of magnitude faster than BLAST, Bioinformatics 26, 2460–2461, doi; 10.1093/bioinformatics/btq461

Edgar, R.C., 2013. UPARSE: highly accurate OTU sequences from microbial amplicon reads. Nature Methods 10, 996–998. doi:10.1038/nmeth.2604

Edgar, R.C., 2016. SINTAX: a simple non-Bayesian taxonomy classifier for 16S and ITS sequences. bioRxiv 074161; doi:10.1101/074161

Erisman, J.W., Galloway, J.N., Seitzinger, S., Bleeker, A., Dise, N.B., Roxana Petrescu, A.M., Leach, A.M., de Vries, W., 2013. Consequences of human modification of the global nitrogen cycle. Philosophical Transactions of the Royal Society B: Biological Sciences 368. doi:10.1098/rstb.2013.0116

Fan, X., Yin, C., Chen, H., Ye, M., Zhao, Y., Li, T., Wakelin, S.A., Liang, Y., 2019. The efficacy of 3,4-dimethylpyrazole phosphate on N2O emissions is linked to niche differentiation of ammonia oxidizing archaea and bacteria across four arable soils. Soil Biology and Biochemistry 130, 82–93.

García-Gil, J.C., Plaza, C., Soler-Rovira, P., Polo, A., n.d. Long-term effects of municipal solid waste compost application on soil enzyme activities and microbial biomass.

Gilsanz, C., Báez, D., Misselbrook, T.H., Dhanoa, M.S., Cárdenas, L.M., 2016. Development of emission factors and efficiency of two nitrification inhibitors, DCD and DMPP. Agriculture, Ecosystems & Environment 216, 1–8. 10.1016/j.agee.2015.09.030

Guo, T., Bai, S.H., Omidvar, N., Wang, Y., Chen, F., Zhang, M., 2023. Insight into the functional mechanisms of nitrogen-cycling inhibitors in decreasing yield-scaled ammonia volatilization and nitrous oxide emission: A global meta-analysis. Chemosphere 338, 139611. doi:10.1016/J.CHEMOSPHERE.2023.139611

Guyomard, H., Bureau J.-C. et al. (2020), Research for AGRI Committee – The Green Deal and the CAP: policy implications to adapt farming practices and to preserve the EU’s natural resources. European Parliament, Policy Department for Structural and Cohesion Policies, Brussels.

Heuermann, D., Gentsch, N., Boy, J., Schweneker, D., Feuerstein, U., Groß, J., Bauer, B., Guggenberger, G., von Wirén, N., 2019. Interspecific competition among catch crops modifies vertical root biomass distribution and nitrate scavenging in soils. Scientific Reports 9, 1–11. doi:10.1038/s41598-019-48060-0

Huérfano, X., Estavillo, J.M., Torralbo, F., Vega-Mas, I., González-Murua, C., Fuertes-Mendizábal, T., 2022. Dimethylpyrazole-based nitrification inhibitors have a dual role in N2O emissions mitigation in forage systems under Atlantic climate conditions. Science of The Total Environment 807, 150670. doi:10.1016/J.SCITOTENV.2021.150670

Kalbitz, K., Solinger, S., Park, J.H., Michalzik, B., Matzner, E., 2000. Controls on the dynamics of dissolved organic matter in soils: A review. Soil Science 165, 277–304. doi:10.109700010694-200004000-00001

Kleineidam, K., Košmrlj, K., Kublik, S., Palmer, I., Pfab, H., Ruser, R., Fiedler, S., Schloter, M., 2011. Influence of the nitrification inhibitor 3,4-dimethylpyrazole phosphate (DMPP) on ammonia-oxidizing bacteria and archaea in rhizosphere and bulk soil. Chemosphere 84, 182–186. doi:10.1016/J.CHEMOSPHERE.2011.02.086

Lazcano, C., Gómez-Brandón, M., Revilla, P., Domínguez, J., 2013. Short-term effects of organic and inorganic fertilizers on soil microbial community structure and function. Biology and Fertility of Soils 49, 723–733. doi:10.1007/s00374-012-0761-7

Lei, J., Fan, Q., Yu, J., Ma, Y., Yin, J., Liu, R., 2022. A meta-analysis to examine whether nitrification inhibitors work through selectively inhibiting ammonia-oxidizing bacteria. Frontiers in Microbiology 13, 962146. doi:10.3389/FMICB.2022.962146/BIBTEX

Li, H. Ruo, Song, X. Tong, Bakken, L.R, Ju, X. Tang, 2023. Reduction of N2O emissions by DMPP depends on the interactions of nitrogen sources (digestate vs. urea) with soil properties. Journal of Integrative Agriculture 22, 251–264. doi:10.1016/J.JIA.2022.08.009

Li, H., Song, X., Wu, D., Wei, D., Li, Y., Ju, X., 2024. Partial substitution of manure increases N2O emissions in the alkaline soil but not acidic soils. Journal of Environmental Management 359, 120993. doi:10.1016/J.JENVMAN.2024.120993

Lognoul, M., Debacq, A., De Ligne, A., Dumont, B., Manise, T., Bodson, B., Heinesch, B., Aubinet, M., 2019. N_2_O flux short-term response to temperature and topsoil disturbance in a fertilized crop: An eddy covariance campaign. Agricultural and Forest Meteorology 271, 193–206. doi: 10.1016/j.agrformet.2019.02.033.

Lori, M., Armengot, L., Schneider, M., Schneidewind, U., Bodenhausen, N., Mäder, P., Krause, H.-M. 2022. Organic management enhances soil quality and drives microbial community diversity in cocoa production systems. Science of The Total Environment 834, 155223. doi:10.1016/j.scitotenv.2022.155223

Lori, M., Hartmann, M., Kundel, D., Mayer, J., Mueller, R.C., Mäder, P., Krause, H.M., 2023. Soil microbial communities are sensitive to differences in fertilization intensity in organic and conventional farming systems, FEMS Microbiology Ecology 99, fiad046. doi:10.1093/femsec/fiad046

Lynch, J., Cain, M., Frame, D., Pierrehumbert, R., 2021. Agriculture’s Contribution to Climate Change and Role in Mitigation Is Distinct From Predominantly Fossil CO_2_-Emitting Sectors. Frontiers in Sustainable Food Systems 4. doi:10.3389/fsufs.2020.518039

Ma, Y., Kang, L., Li, Y., Zhang, X., Cardenas, L.M., Chen, Q., 2023. Is sorghum a promising summer catch crop for reducing nitrate accumulation and enhancing eggplant yield in intensive greenhouse vegetable systems? Plant and Soil 499, 1–13. doi:10.1007/S11104-023-05923-W

McMurdie, P.J., Holmes, S., 2013. phyloseq: an R package for reproducible interactive analysis and graphics of microbiome census data. PLoS One. 8, e61217. doi:10.1371/journal.pone.0061217

Mooshammer, M., Wanek, W., Hämmerle, I., Fuchslueger, L., Hofhansl, F., Knoltsch, A., et al., 2014. Adjustment of microbial nitrogen use efficiency to carbon:nitrogen imbalances regulates soil nitrogen cycling. Nature Communications 5, 3694. doi: 10.1038/ncomms4694

O’Sullivan, C.A., Fillery, I.R.P., Roper, M.M., Richards, R.A., 2016. Identification of several wheat landraces with biological nitrification inhibition capacity. Plant and Soil 404, 61–74. doi:10.1007/s11104-016-2822-4

Oksanen, J., Blanchet, F.G., Friendly, M., Kindt, R., Legendre, P., McGlinn, D., Minchin, P.R., O’Hara, R.B., Simpson, G.L., Solymos, P., Stevens, M.H.H., Szoecs, E., Wagner, H. 2020. Vegan: Community Ecology Package. R package version 2.5–7. https://CRAN.R-project.org/package=vegan

Otaka, J., Subbarao, G.V., Ono, H., Yoshihashi, T., 2021. Biological nitrification inhibition in maize— isolation and identification of hydrophobic inhibitors from root exudates. Biology and Fertility of Soils 1–14. doi:10.1007/S00374-021-01577-X/FIGURES/10

Ouyang, Y., Norton, J.M., Stark, J.M., 2017. Ammonium availability and temperature control contributions of ammonia oxidizing bacteria and archaea to nitrification in an agricultural soil. Soil Biology and Biochemistry 113, 161–172.

Peixoto, L., Petersen, S.O., 2023. Efficacy of three nitrification inhibitors to reduce nitrous oxide emissions from pig slurry and mineral fertilizers applied to spring barley and winter wheat in Denmark. Geoderma Regional 32, e00597. doi:10.1016/J.GEODRS.2022.E00597

Petersen, S.O., Peixoto, L.E.K., Sørensen, H., Tariq, A., Brændholt, A., Hansen, L.V., Abalos, D., Christensen, A.T., Nielsen, C.S., Pullens, J.W.M., Bruun, S., Jensen, L.S., Olesen, J.E., 2023. Higher N2O emissions from organic compared to synthetic N fertilisers on sandy soils in a cool temperate climate. Agriculture, Ecosystems & Environment 358, 108718. doi:10.1016/J.AGEE.2023.108718

Pilegaard, K., 2013. Processes regulating nitric oxide emissions from soils. Philosophical Transactions of the Royal Society of London. Series B, Biological Sciences 368, 20130126. doi:10.1098/rstb.2013.0126

Quast, C., Pruesse, E., Yilmaz, P., Gerken, J., Schweer, T., Yarza, P., Peplies, J., Glöckner, F.O., 2013. The SILVA ribosomal RNA gene database project: improved data processing and web-based tools. Nucleic Acids Research 41, D590–6. doi:10.1093/nar/gks1219

Sadhukhan, R., Jatav, H.S., Sen, S., Sharma, L.D., Rajput, V.D., Thangjam, R., Devedee, A.K., Singh, S.K., Gorovtsov, A., Choudhury, S., Patra, K., 2022. Biological nitrification inhibition for sustainable crop production. Plant Perspectives to Global Climate Changes: Developing Climate-Resilient Plants 135–150. doi:10.1016/B978-0-323-85665-2.00007-8

Saggar, S., Jha, N., Deslippe, J., Bolan, N.S., Luo, J., Giltrap, D.L., Kim, D.-G., Zaman, M., Tillman, R.W., 2013. Denitrification and N2O:N2 production in temperate grasslands: Processes, measurements, modelling and mitigating negative impacts. Science of The Total Environment 465, 173–195. 10.1016/j.scitotenv.2012.11.050

Saud, S., Wang, D., Fahad, S., 2022. Improved Nitrogen Use Efficiency and Greenhouse Gas Emissions in Agricultural Soils as Producers of Biological Nitrification Inhibitors. Frontiers in Plant Science 13, 534. doi:10.3389/fpls.2022.854195

Schmieder, R., Edwards, R., 2011. Quality control and preprocessing of metagenomic datasets, Bioinformatics 27, 863–864, doi:10.1093/bioinformatics/btr026

Soares, J.R., Souza, B.R., Mazzetto, A.M., Galdos, M. V., Chadwick, D.R., Campbell, E.E., Jaiswal, D., Oliveira, J.C., Monteiro, L.A., Vianna, M.S., Lamparelli, R.A.C., Figueiredo, G.K.D.A., Sheehan, J.J., Lynd, L.R., 2023. Mitigation of nitrous oxide emissions in grazing systems through nitrification inhibitors: a meta-analysis. Nutrient Cycling in Agroecosystems 125, 359–377. doi:10.1007/s10705-022-10256-8

Subbarao, G. V., Kishii, M., Bozal-Leorri, A., Ortiz-Monasterio, I., Gao, X., Ibba, M.I., Karwat, H., Gonzalez-Moro, M.B., Gonzalez-Murua, C., Yoshihashi, T., Tobita, S., Kommerell, V., Braun, H.-J., Iwanaga, M., 2021. Enlisting wild grass genes to combat nitrification in wheat farming: A nature-based solution. Proceedings of the National Academy of Sciences 118, e2106595118. doi:10.1073/PNAS.2106595118

Subbarao, G. V., Searchinger, T.D., 2021. A “more ammonium solution” to mitigate nitrogen pollution and boost crop yields. Proceedings of the National Academy of Sciences of the United States of America. doi:10.1073/pnas.2107576118

Subbarao, G.V., Rondon, M., Ito, O., Ishikawa, T., Rao, I.M., Nakahara, K., Lascano, C., Berry, W.L., 2007a. Biological nitrification inhibition (BNI)—is it a widespread phenomenon? Plant and Soil 294, 5–18. doi:10.1007/s11104-006-9159-3

Subbarao, G.V., Tomohiro, B., Masahiro, K., Osamu, I., Samejima, H., Wang, H.Y., Pearse, S.J., Gopalakrishnan, S., Nakahara, K., Zakir Hossain, A.K.M., Tsujimoto, H., Berry, W.L., 2007b. Can biological nitrification inhibition (BNI) genes from perennial Leymus racemosus (Triticeae) combat nitrification in wheat farming? Plant and Soil 299, 55–64. doi:10.1007/s11104-007-9360-z

Suleiman, A.K.A., Gonzatto, R., Aita, C., Lupatini, M., Jacques, R.J.S., Kuramae, E.E., Antoniolli, Z.I., Roesch, L.F.W., 2016. Temporal variability of soil microbial communities after application of dicyandiamide-treated swine slurry and mineral fertilizers. Soil Biology and Biochemistry 97, 71–82. doi:10.1016/J.SOILBIO.2016.03.002

Sun, H., Li, Youfa, Xing, Y., Bodington, D., Huang, X., Ding, C., Ge, T., Di, H., Xu, J., Gubry-Rangin, C., Li, Yong, 2024. Organic fertilizer significantly mitigates N2O emissions while increase contributed of comammox Nitrospira in paddy soils. Science of the Total Environment 954. doi:10.1016/J.SCITOTENV.2024.176578

Torralbo, F., Menéndez, S., Barrena, I., Estavillo, J.M., Marino, D., González-Murua, C., 2017. Dimethyl pyrazol-based nitrification inhibitors effect on nitrifying and denitrifying bacteria to mitigate N2O emission. Scientific Reports 7, 13810. doi:10.1038/s41598-017-14225-y

Tufail, M.A., Irfan, M., Umar, W., Wakeel, A., Schmitz, R.A., 2023a. Mediation of gaseous emissions and improving plant productivity by DCD and DMPP nitrification inhibitors: Meta-analysis of last three decades. Environmental Science and Pollution Research 30, 64719–64735. doi:10.1007/s11356-023-26318-5

Tufail, M.A., Irfan, M., Umar, W., Wakeel, A., Schmitz, R.A., 2023b. Mediation of gaseous emissions and improving plant productivity by DCD and DMPP nitrification inhibitors: Meta-analysis of last three decades. Environmental Science and Pollution Research 30, 64719–64735. doi:10.1007/S11356-023-26318-5/FIGURES/8

United Nations, 2019. World Population Prospects 2019, Table: Total Population – Both Sexes, Department of Economic and Social Affairs, Population Division, UN, New York. https://population.un.org/wpp/Download/Standard/Population/. [Accessed november 2023]].

Valkama, E., Tzemi, D., Esparza-Robles, U.R., Syp, A., O’Toole, A., Maenhout, P., 2024. Effectiveness of soil management strategies for mitigation of N2O emissions in European arable land: A meta-analysis. European Journal of Soil Science 75, e13488. doi:10.1111/EJSS.13488

Vega-Mas, I., Ascencio-Medina, E., Menéndez, S., González-Torralba, J., González-Murua, C., Marino, D., González-Moro, M.B., 2024. Selecting an optimal sorghum cultivar can improve nitrogen availability and wheat yield in crop rotation. Journal of the Science of Food and Agriculture. doi:10.1002/JSFA.13969

Wang, Q., Zhao, Z., Yuan, M., Zhang, Z., Chen, S., Ruan, Y., Huang, Q., 2022. Impacts of urea and 3,4-dimethylpyrazole phosphate on nitrification, targeted ammonia oxidizers, non-targeted nitrite oxidizers, and bacteria in two contrasting soils. Frontiers in Microbiology 13. doi:10.3389/FMICB.2022.952967

Wang, X., Bai, J., Xie, T., Wang, W., Zhang, G., Yin, S., Wang, D., 2021. Effects of biological nitrification inhibitors on nitrogen use efficiency and greenhouse gas emissions in agricultural soils: A review. Ecotoxicology and Environmental Safety. doi:10.1016/j.ecoenv.2021.112338

Ward, B.B., 2015. Nitrification, in: Encyclopedia of Ecology: Volume 1-4, Second Edition. Elsevier, pp. 351–358. doi:10.1016/B978-0-12-409548-9.00697-7

Wu, D., Zhang, Y., Dong, G., Du, Z., Wu, W., Chadwick, D., Bol, R., 2021. The importance of ammonia volatilization in estimating the efficacy of nitrification inhibitors to reduce N2O emissions: A global meta-analysis. Environmental Pollution 271, 116365. doi:10.1016/J.ENVPOL.2020.116365

Wu, G., Yang, S., Luan, C. sheng, Wu, Q., Lin, L. li, Li, X. xiao, Che, Z., Zhou, D. bao, Dong, Z. rong, Song, H., 2024. Partial organic substitution for synthetic fertilizer improves soil fertility and crop yields while mitigating N2O emissions in wheat-maize rotation system. European Journal of Agronomy 154, 127077. doi:10.1016/J.EJA.2023.127077

Yan, X., Gong, W., 2010. The role of chemical and organic fertilizers on yield, yield variability and carbon sequestration— results of a 19-year experiment. Plant and Soil 331, 471–480. doi:10.1007/s11104-009-0268-7

Yang, M., Fang, Y., Sun, D., Shi, Y., 2016a. Efficiency of two nitrification inhibitors (dicyandiamide and 3, 4-dimethypyrazole phosphate) on soil nitrogen transformations and plant productivity: A meta-analysis. Scientific Reports 6, 22075. doi:10.1038/srep22075

Yang, M., Fang, Y., Sun, D., Shi, Y., 2016b. Efficiency of two nitrification inhibitors (dicyandiamide and 3, 4-dimethypyrazole phosphate) on soil nitrogen transformations and plant productivity: a meta-analysis. Scientific Reports 2016 6:1 6, 1–10. doi:10.1038/srep22075

Yin, M., Gao, X., Kuang, W., Zhang, Y., 2023. Meta-analysis of the effect of nitrification inhibitors on the abundance and community structure of N2O-related functional genes in agricultural soils. Science of the Total Environment 865. doi:10.1016/J.SCITOTENV.2022.161215

Yuan, Y., Wang, M., Feng, X., Li, Q., Qin, Y., Sun, B., Li, C., Zhang, J., Liu, H. 2023. Effects of Nitrogen Fertilizer on *Nitrospira*- and *Nitrobacter*-like Nitrite-Oxidizing Bacterial Microbial Communities under Mulched Fertigation System in Semi-Arid Area of Northeast China. Agronomy 13, 2909. doi:10.3390/agronomy13122909

Zhang, M., Gao, X., Chen, G., Afzal, M.R., Wei, T., Zeng, H., Subbarao, G. V., Wei, Z., Zhu, Y., 2023. Intercropping with BNI-sorghum benefits neighbouring maize productivity and mitigates soil nitrification and N2O emission. Agriculture, Ecosystems and Environment 352, 108510. doi:10.1016/J.AGEE.2023.108510

Zhang, M., Wang, W., Zhang, Y., Teng, Y., Xu, Z., 2017. Effects of fungicide iprodione and nitrification inhibitor 3, 4-dimethylpyrazole phosphate on soil enzyme and bacterial properties. Science of The Total Environment 599–600, 254–263. doi:10.1016/J.SCITOTENV.2017.05.011

Zhang, Y., Wang, W., Yao, H., 2022. Urea-based nitrogen fertilization in agriculture: a key source of N_2_O emissions and recent development in mitigating strategies. Archives of Agronomy and Soil Science. doi:10.1080/03650340.2022.2025588

Zhou, J., Guan, D., Zhou, B., Zhao, B., Ma, M., Qin, J., Jiang, X., Chen, S., Cao, F., Shen, D., Li, J., 2015. Influence of 34-years of fertilization on bacterial communities in an intensively cultivated black soil in northeast China. Soil Biology and Biochemistry 90, 42–51. doi:10.1016/J.SOILBIO.2015.07.005

Zhou, X., Wang, S., Ma, S., Zheng, X., Wang, Z., Lu, C., 2020. Effects of commonly used nitrification inhibitors—dicyandiamide (DCD), 3,4-dimethylpyrazole phosphate (DMPP), and nitrapyrin—on soil nitrogen dynamics and nitrifiers in three typical paddy soils. Geoderma 380, 114637. doi:10.1016/j.geoderma.2020.114637

