## Appendix A. Supplementary data. for "From Synthetic to Biological Nitrification Inhibition: Advancing Stabilization of Organic Fertilizers"

### Supplementary material:

**Supplementary Table S1.** Chemical properties of the soil analyzed at Laboratoires d’analyses agricoles, La Hulpe, Belgium. Sieved and homogenized soil was used as a single pool to set up the experiments. For EXP2, soil (mixed with sand at 2:1, w:w) collected from pots was analyzed before initiating the incubations. NT, not treated soil; SDMPP, soil treated with DMPP; P44, rhizospheric soil from wheat cv. Persia 44; PP, rizhospheric soil from white mustard cv. Pole Position (PP) and VERDI, rizhospheric soil from white mustard cv. Verdi.

| | **Soil chemical properties** | | | | | | | **Soil texture:** *Silt loam* | |
|---|---|---|---|---|---|---|---|---|---|
| | **Soil EXP 1** | **Soil:sand (2:1, w:w) EXP 2** | | | | | | **EXP 1** | **EXP 2** |
| | | NT | SDMPP | Rhizospheric soil | | | | | |
| | | | | P44 | PP | Verdi | | | |
| **pH**(KCl) | 7.4 | 6.2 | 6.5 | 6.2 | 6.2 | 6.2 | Sand (%) | 10.3 | 8.3 |
| **Organic matter** (% of dry soil) | 1.1 | 9 | 9 | 6 | 9 | 11 | Silt (%) | 76.2 | 77.6 |
| **Total nitrogen** (% of dry soil) | 0.11 | 0.09 | 0.1 | <0.06 | 0.09 | 0.11 | Clay (%) | 13.5 | 13.5 |
| **C/N ratio** | 10 | 10 | 10 | 10 | 10 | 10 | | | |

**Supplementary Table S2.** Classification of fertilization treatments, and percentage of N in form of ammonium N-$NH_4^+$(%), pH value, C/N ratio and dry matter content (%) of each mineral or organic fertilizer.

| **ID** | **Descripción** | **Fertilizer form** | **Origin** | **Dry Matter** (% of fresh product) | **pH** | **C/N ratio** | **Total N** (%) | **N-$NH_4^+$** (%) |
|---|---|---|---|---|---|---|---|---|
| | **EXP1** | | | | | | | |
| **D** | Digestate | Liquid | Derived from plant waste | 6.5 | 8.1 | 4.5 | 6.7 | 46 |
| **O7** | Orgamine 7 | Solid | Plant-animal mixed | 92.9 | 9.1 | 3.1 | 10.2 | 31 |
| **CM** | Chicken manure | Solid | Animal. Mixed with straw | 55.4 | 6.5 | 8.7 | 5.2 | 13 |
| **CS** | Cattle slurry | Liquid | Animal. | 8.2 | 6.8 | 10.4 | 4.2 | 31 |
| **AS** | Ammonium sulfate | Control mineral fertilizer | | | | | | 100 |
| **NF** | Non-fertilized | | | | | | | |
| | **EXP2** | | | | | | | |
| **D** | Digestate | Liquid | Derived from plant waste | 8.0 | 8.4 | 7.4 | 2.9 | 48 |
| **O7** | Orgamine 7 | Solid | Plant-animal mixed | 93.0 | 9.1 | 3.1 | 10.2 | 31 |

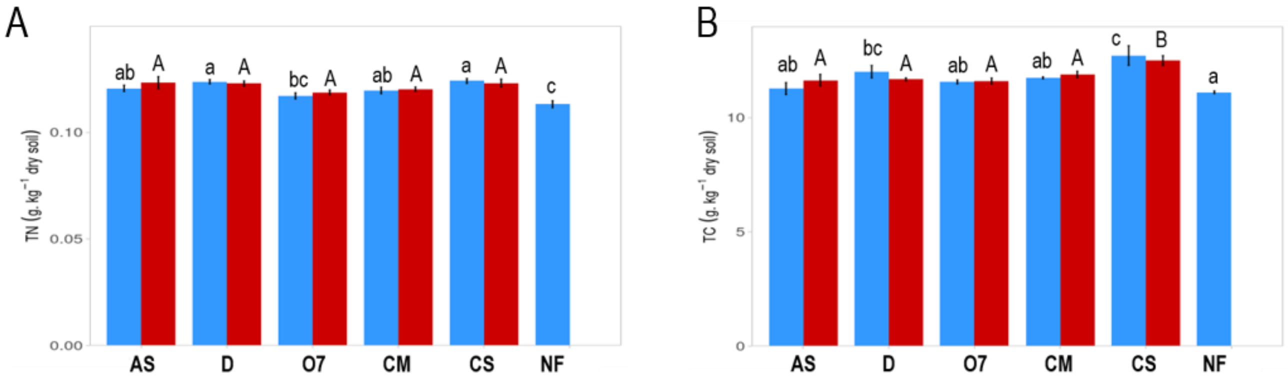


**Supplementary Fig. S1.** Total carbon (TC) and total nitrogen (TN) contents in soil-fertilizer mixture samples collected at Day 0 (EXP1). Values represent mean ± SE (n = 4). Letters represent significant differences between treatments analyzed by Duncan's test (P<0.05), lower case letters for treatments without the inhibitor and upper case letters for treatments with DMPP.

**Supplementary Text 1:** Rhizospheric soil collection and incubation set up for EXP2.

About 20 seeds of each selected genotype (P44, PP and VERDI) were first germinated in Petri dishes containing a wet filter paper. The Petri dishes were kept in cold room (4°C) in the dark for 48h to break dormancy, then transfer into an incubator at 20°C for 24 h. Germinated seeds were then transplanted to pots (10x10x10cm) containing the equivalent of 1 kg DW soil. This silty soil was collected from a field in Gembloux (50°33’36.0"N 4°42’36.0"E) (**Table S1)**. The soil was air-dried for four days and then homogenized, sieved at 3.5 mm and kept at 4 °C until use. Soil was then mixed with sand (2:1, w:w) to enhance porosity and ensure the proper development of plant roots. Four pots (1L) per genotype were prepared, each containing the equivalent of 1 kg DW substrate and three plants per pot. Pots were watered to reach 60% water-filled pore space (WFPS) to promote nitrification process and soil moisture was adjusted every 2-3 days. Pots were placed in a growth chamber at 20°C for four weeks. Each pot was fertilized once a week with ammonium sulphate (AS) at a rate of 20 mg N $kg^{-1}$ DW soil. After four weeks of growth, the shoot was removed and the soil was collected. Roots were removed as much as possible, and the rhizospheric soil from the four pots of each genotype was thoroughly mixed for homogenization before being packed into 250 mL glass jars for incubation. Subsamples of each rhizospheric soil were sent to Laboratoires d’Analyses Agricoles, La Hulpe, Belgium, for the analyses presented in **Table S1.**

During crop growth, eight additional pots containing 1 kg equivalent DW soil were placed in the same growth chamber, fertilized and watered similarly to the plant-containing pots. In four of the pots, DMPP was added to the fertilizer at 1% of the supplied N (SDMPP treatment). As for rhizospheric soil, after four weeks in the growth chamber and before packing the soil into incubation bottles, subsamples of standard soil and soil treated with DMPP were collected and sent to Laboratoires d’Analyses Agricoles, La Hulpe, Belgium for analyses presented in **Table S1.**

**Supplementary Table S3.** Primers sequences used in qPCR for the quantification of bacterial and archaeal marker genes (From Krause et al., 2017 and Torralbo et al., 20017).

| Marker gene | ID | Primer sequence (5' – 3') |
|---|---|---|
| Bacterial *16S rRNA*: | 341F<br>534R | CCTACGGGAGGCAGCAG<br>ATTACCGCGGCTGCTGGCA |
| Archaeal *16S rRNA*: | 771F<br>957R | ACGGTGAGGGATGAAAGCT<br>CGGCGTTGACTCCAATTG |
| bacterial *amoA:* | amoA1F<br>amoA2R | GGGGTTTCTACTGGTGGT<br>CCCCTCKGSAAAGCCTTCTTC |
| archaeal *amoA*: | amo19F<br>CrenamoA616r48x R | ATGGTCTGGCTWAGACG<br>GCCATCCABCKRTANGTCCA |
| *nirS*: | nirScd3af<br>R3cd | AACGYSAAGGARACSGG<br>GASTTCGGRTGSGTCTTGA |
| *nirK*: | nirK876C<br>nirK1040 | ATYGGCGGVCAYGGCGA<br>GCCTCGATCAGRTTRTGGTT |
| *nosZ:* | nosZ2F<br>nosZ2R | CGCRACGGCAASAAGGTSMSSGT<br>CAKRTGCAKSGCRTGGCAGAA |
| *nosZ-II*: | nosZ-II-F<br>nosZ-II-R | CTIGGICCIYTKCAYAC<br>GCIGARCARAAITCBGTRC |

**Supplementary Table S4.** Details for qPCR reactions and reaction mixtures used for the quantification of bacterial and archaeal marker genes (Modified from Krause et al., 2017 and Torralbo et al., 20017).

| Target gene | qPCR conditions | Reaction mixtures | Volumes [µl] |
|---|---|---|---|
| *Bacterial 16S rRNA* | 95ºC – 2 min<br><br>95ºC – 15 sec<br>**60** °C – 30 sec<br>72 °C – 30 sec<br>80 °C – 30sec<br>x 40 cycles | Kapa SYBR® (2X)<br>Primer F (5 µM)<br>Primer R (5 µM)<br>PCR water<br>DNA – template | 5<br>1<br>1<br>2<br>1 |
| *Archaeal 16S rRNA* | 95ºC – 2 min<br><br>95ºC – 15 sec<br>**58** °C – 30 sec<br>72 °C – 30 sec<br>80 °C – 30sec<br>x 40 cycles | Kapa SYBR® (2X)<br>Primer F (5 µM)<br>Primer R (5 µM)<br>PCR water<br>DNA – template | 5<br>1<br>1<br>2<br>1 |
| AOB | 94°C – 3 min<br><br>94°C – 30 sec<br>**59.5**°C – 30 sec<br>68°C – 30 sec<br>x 40 cycles<br><br>68°C – 5 min | Kapa SYBR® (2X)<br>Primer F (5 µM)<br>Primer R (5 µM)<br>BSA (2% w/v)<br>PCR water<br>DNA – template | 5<br>0.75<br>0.75<br>2<br>0.5<br>1 |
| AOA | 94°C – 3 min<br><br>94°C – 30 sec<br>**55**°C – 30 sec<br>68°C – 30 sec<br>x 40 cycles<br><br>68°C – 5 min | Kapa SYBR® (2X)<br>Primer F (5 µM)<br>Primer R (5 µM)<br>PCR water<br>DNA – template | 5<br>1<br>1<br>2<br>1 |
| *nirS* | 94°C – 3 min<br><br>94°C – 30 sec<br>**58**°C – 30 sec<br>68°C – 30 sec<br>x 40 cycles<br><br>68°C – 5 min | Kapa SYBR® (2X)<br>Primer F (5 µM)<br>Primer R (5 µM)<br>PCR water<br>BSA<br>DNA - template | 5<br>1<br>1<br>1.6<br>0.4<br>1 |
| *nirK* | 94°C – 3 min<br><br>94°C – 15 sec<br>**62-57**°C – 20 sec<br>68°C – 30 sec<br>x 6 cycles<br><br>94°C – 30 sec<br>**57**°C – 30 sec<br>68°C – 30 sec<br>x 35 cycles<br><br>68°C – 5 min | Kapa SYBR® (2X)<br>Primer F (2 µM)<br>Primer R (2 µM)<br>PCR water<br>DNA – template | 5<br>1<br>1<br>2<br>1 |
| *nosZ* | 94°C – 3 min<br><br>94°C – 15 sec<br>**65-60**°C – 20 sec (gradient)<br>68°C – 30 sec<br>x 6 cycles<br><br>94°C – 30 sec<br>**60**°C – 30 sec<br>68°C – 30 sec<br>x 35 cycles<br><br>68°C – 5 min | Kapa SYBR® (2X)<br>Primer F (5 µM)<br>Primer R (5 µM)<br>PCR water<br>DNA – template | 5<br>1<br>1<br>2<br>1 |
| nosZ-II | 94°C – 3 min<br><br>94°C – 30 sec<br>**54**°C – 30 sec<br>68°C – 30 sec<br>x 40 cycles<br><br>68°C – 5 min | Kapa SYBR® (2X)<br>Primer F (10 µM)<br>Primer R (10 µM)<br>DMSO<br>PCR water<br>DNA – template | 5<br>1<br>1<br>0.4<br>1.6<br>1 |

**Supplementary Table S5.** Standard plasmids and genes

| **Target gene** | **Plasmid** | **Source of standard** | **Size of insert (pb)** | **Size of apmplicon (pb)** |
|---|---|---|---|---|
| *Bacterial 16S rRNA* | p-GEMT | *Pseudomonas aeruginosa PAO1* | 194 | 194 |
| *Archaeal 16S rRNA* | p-GEMT | *Fosmid clone 29i4* | 257 | 257 |
| *AOB* | pCR2.1 | *Synthetic from Nitrosomonas eutropha C91* | 491 | 491 |
| *AOA* | pCR2.1 | *Synthetic from fosmid clone 54d9* | 628 | 628 |
| *nirS* | pCR4-TOPO | *Ralstonia eutropha H16* | 1607 | 413 |
| *nirK* | pCR4-TOPO | *Ensifer meliloti 1021* | 542 | 162 |
| *nosZ* | pCR4-TOPO | *Ensifer meliloti 1021* | 1884 | 267 |
| *nosZ-II* | pEX-A | *Gemmatimonas aurantiaca* | 800 | 690-720 |

**Supplementary Table S6.** Amount of sequences and OTUs before and after filtering non-bacterial taxa

| | Sequences | OTUs |
|---|---|---|
| Raw | 4058775 | 6802 |
| Cyanobacteria | 15221 | 99 |
| Mitochondria | 89713 | 377 |
| Archaea | 79052 | 36 |
| Filtered | 3874789 | 6290 |

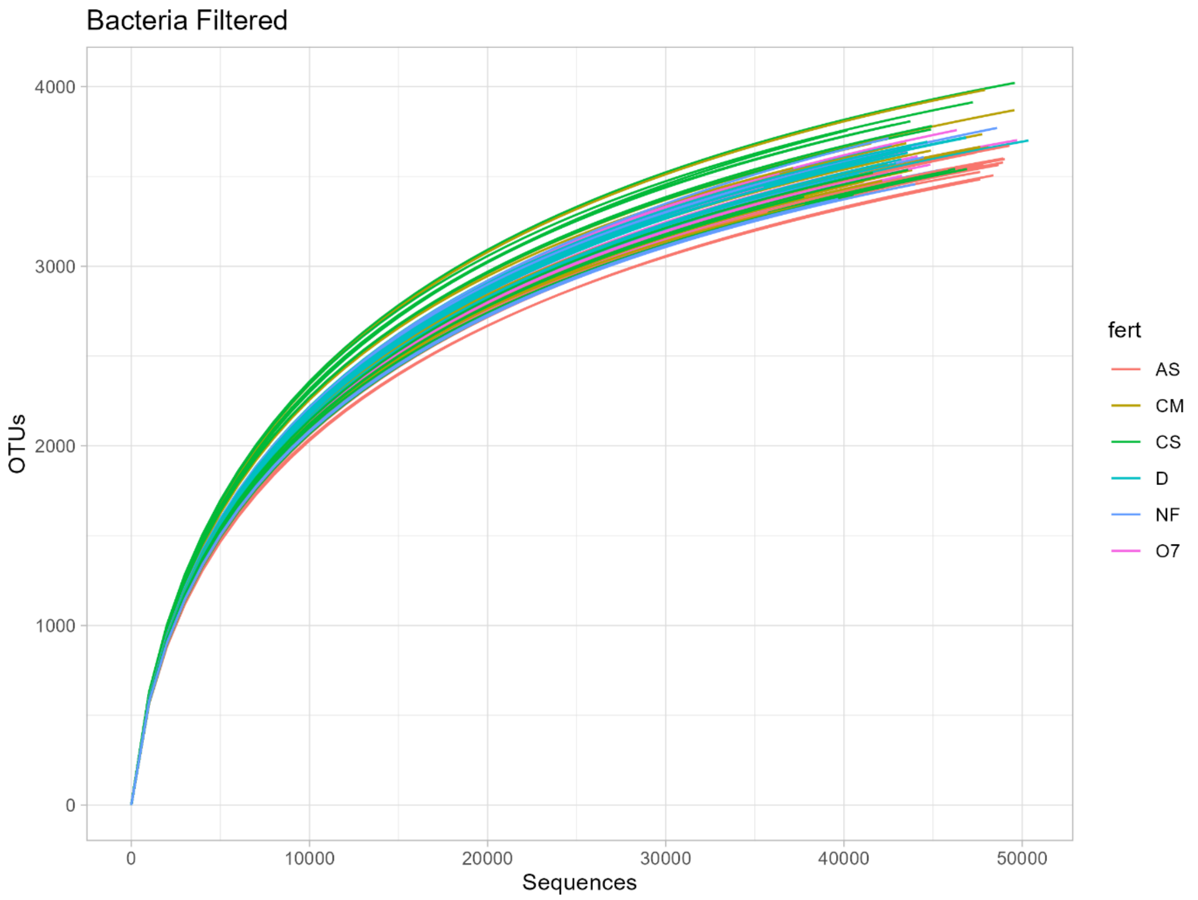


**Supplementary Figure S2.** Rarefaction curves of bacterial sequences for each sample, color-grouped according to fertilization treatment (n=88). Amount of sequences per sample range from 34608 to 50457. D, liquid digestate; O7, Orgamine 7 pellets; CM, chicken manure; CS, cattle slurry (CS); NF, non-fertilized control.

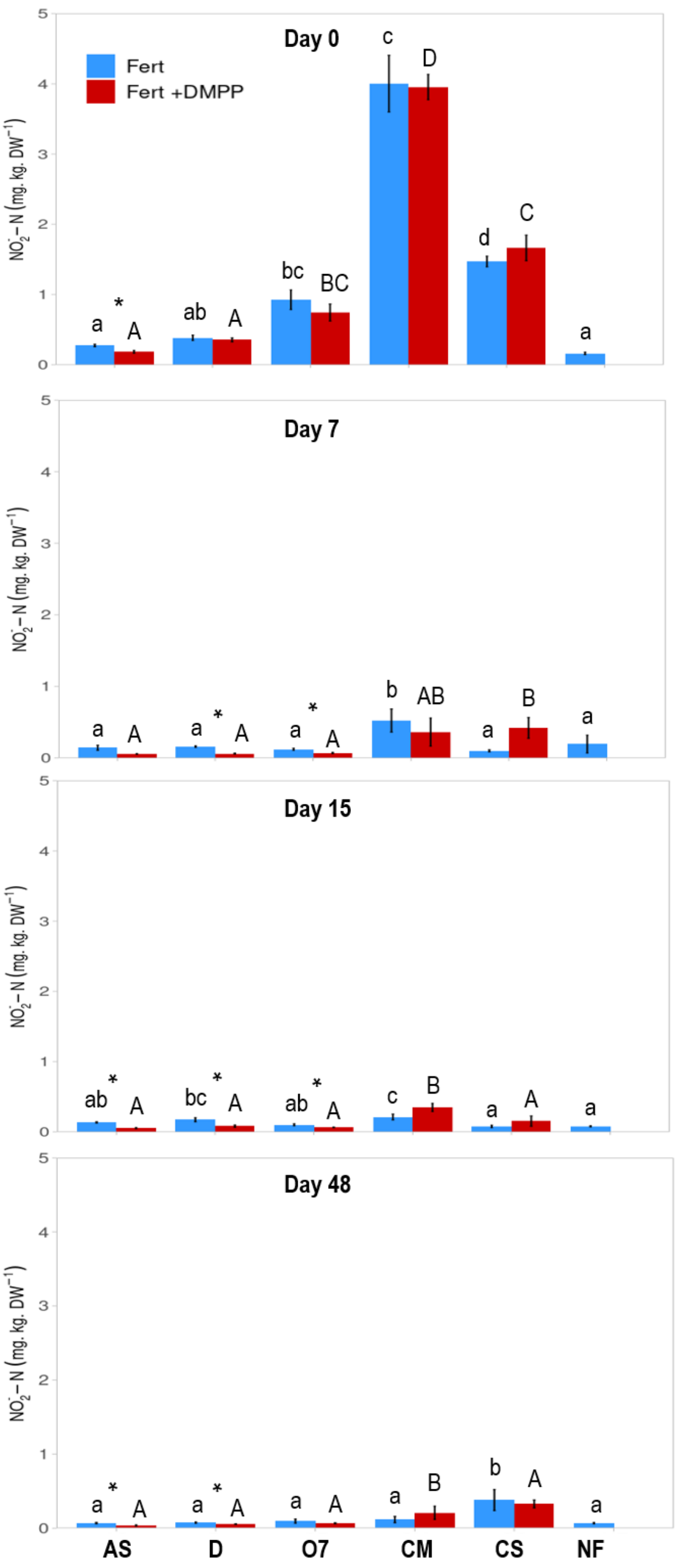


**Supplementary Fig. S3.** Soil nitrite ($NO_2^-$-N) content for each fertilization treatment without (blue) and with the addition of DMPP (red). Values represent mean ± SE (n = 4). Asterisk (*) indicates significant differences between the same fertilizer with and without DMPP (T-student, $p < 0.05$). Letters represent significant differences between treatments analyzed by Duncan's test ($P<0.05$), lower case letters for treatments without the inhibitor and upper case letters for treatments with DMPP. D, liquid digestate; O7, Orgamine 7 pellets; CM, chicken manure; CS, cattle slurry (CS); NF, non-fertilized control.

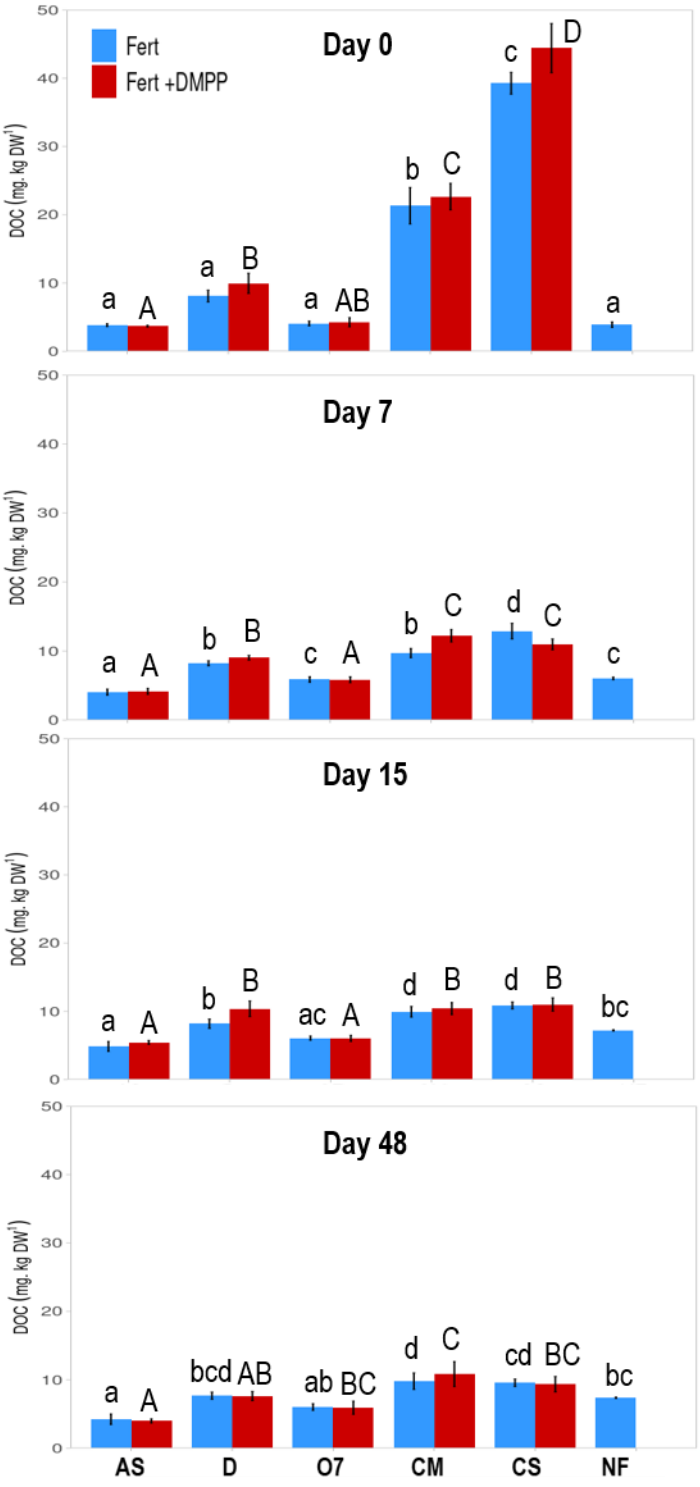


**Supplementary Fig. S4.** Soil DOC value for each fertilization treatment without (blue) and with the addition of DMPP (red). Values represent mean ± SE (n = 4). Statistics as in Supplementary Fig. S3.

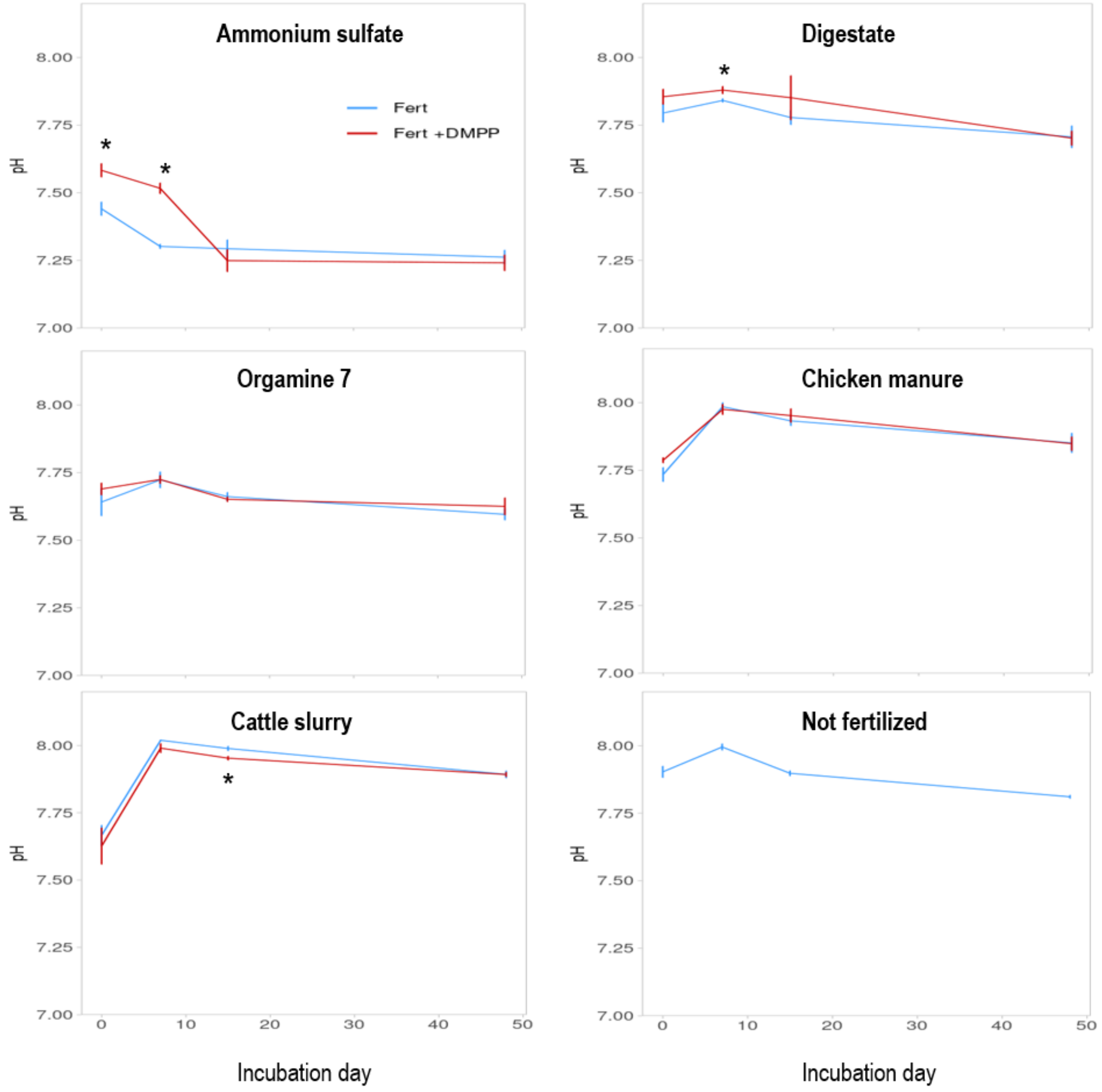


**Supplementary Fig. S5.** Soil pH value for each fertilization treatment without (blue lines) and with the addition of DMPP (red slines). Values represent mean ± SE (n = 4). Asterisk (*) indicates significant differences between same fertilizer with and without DMPP (T-student, $p < 0.05$).

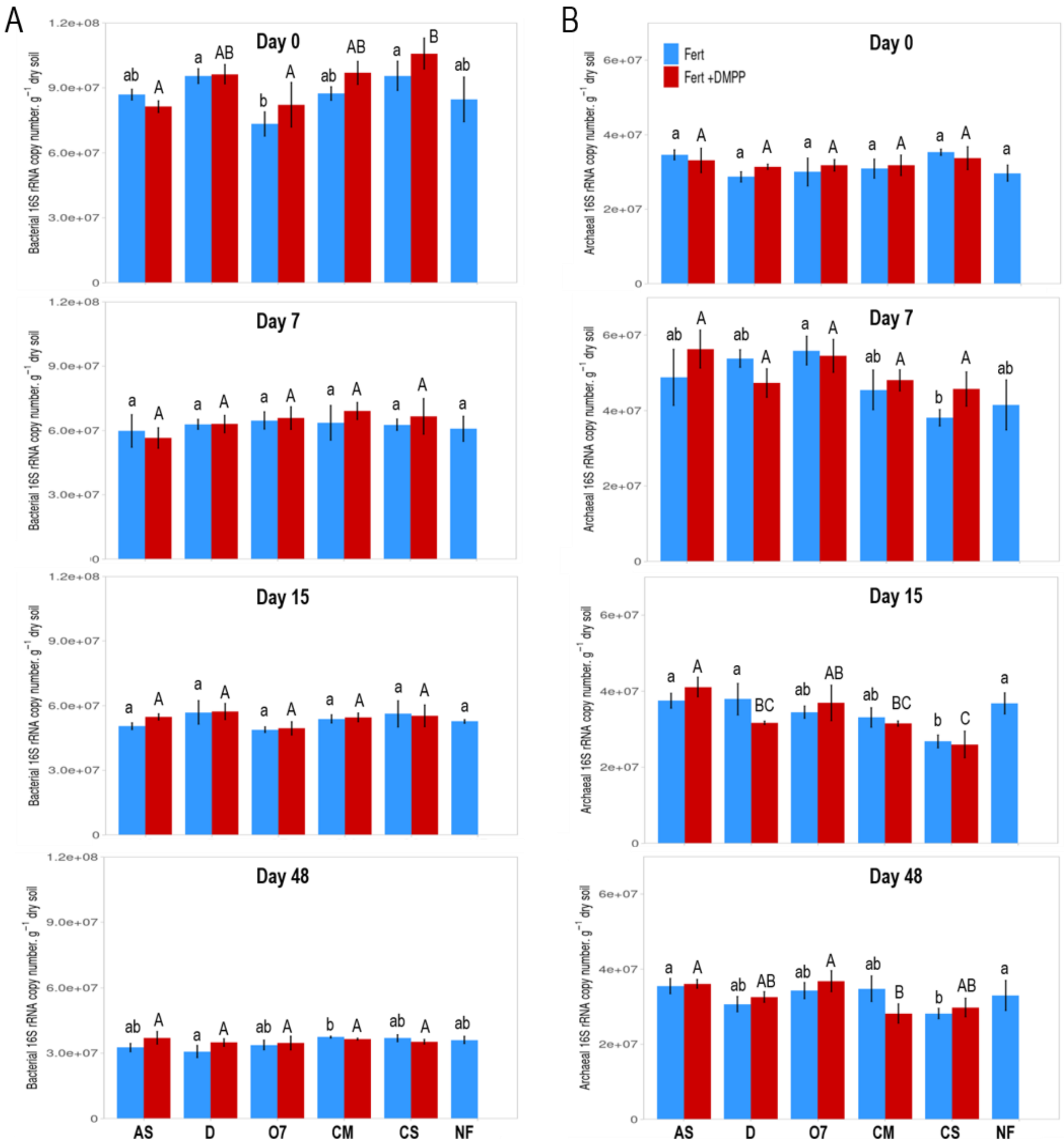

**Supplementary Fig. S6.** Abundance of total bacterial (A) and archaeal (B) population determined by 16Sr RNA marker gene at Day 0, 7, 15 and 48 after fertilization. Values represent mean ± SE (n = 4). Statistics as in Supplementary Fig. S3.

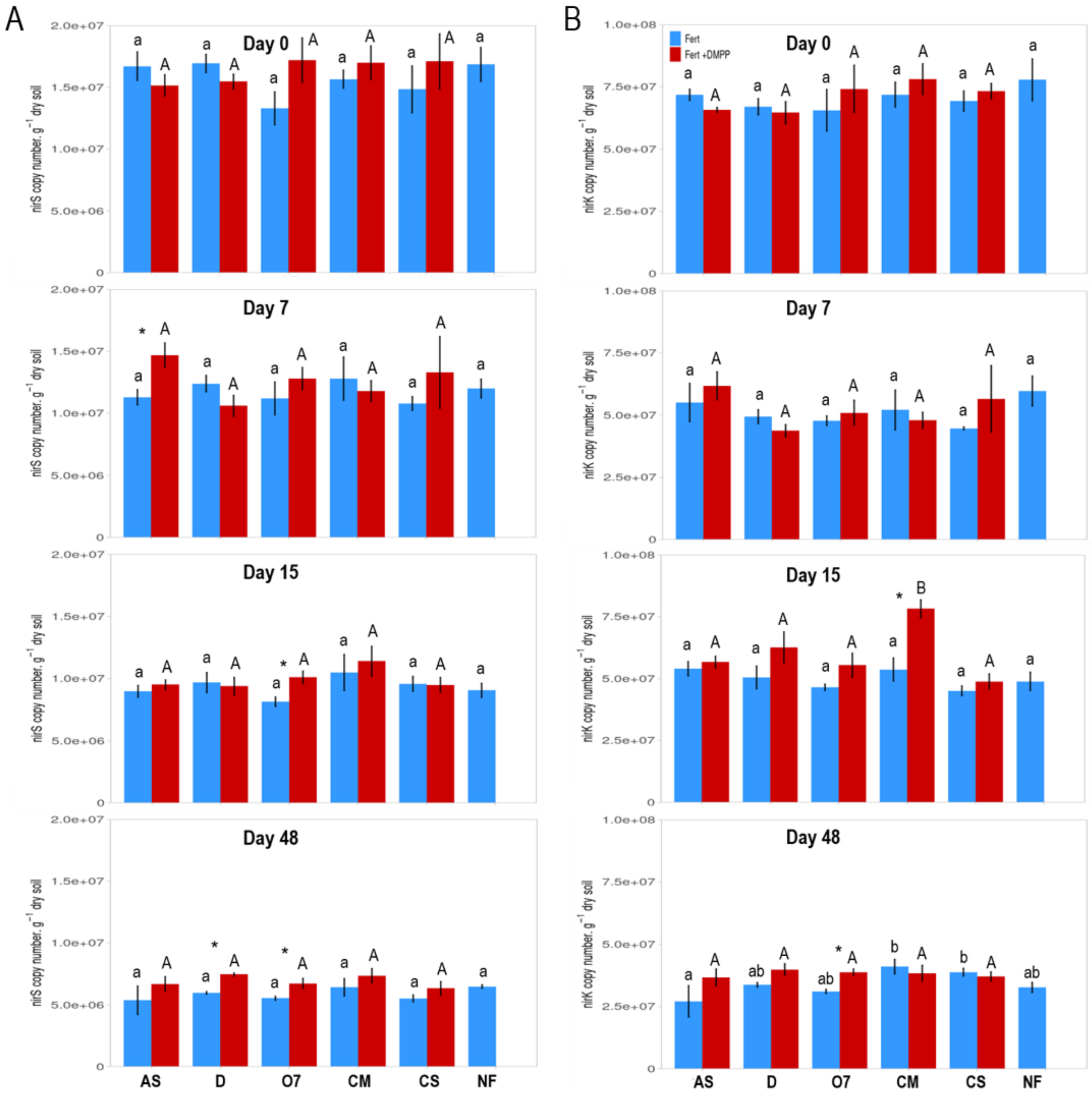


**Supplementary Fig. S7.** Abundance of total marker genes *nirK* and *nirS* determined at Day 0, 7, 15 and 48 after fertilization. Values represent mean ± SE (n = 4). Statistics as in Supplementary Fig. S3.

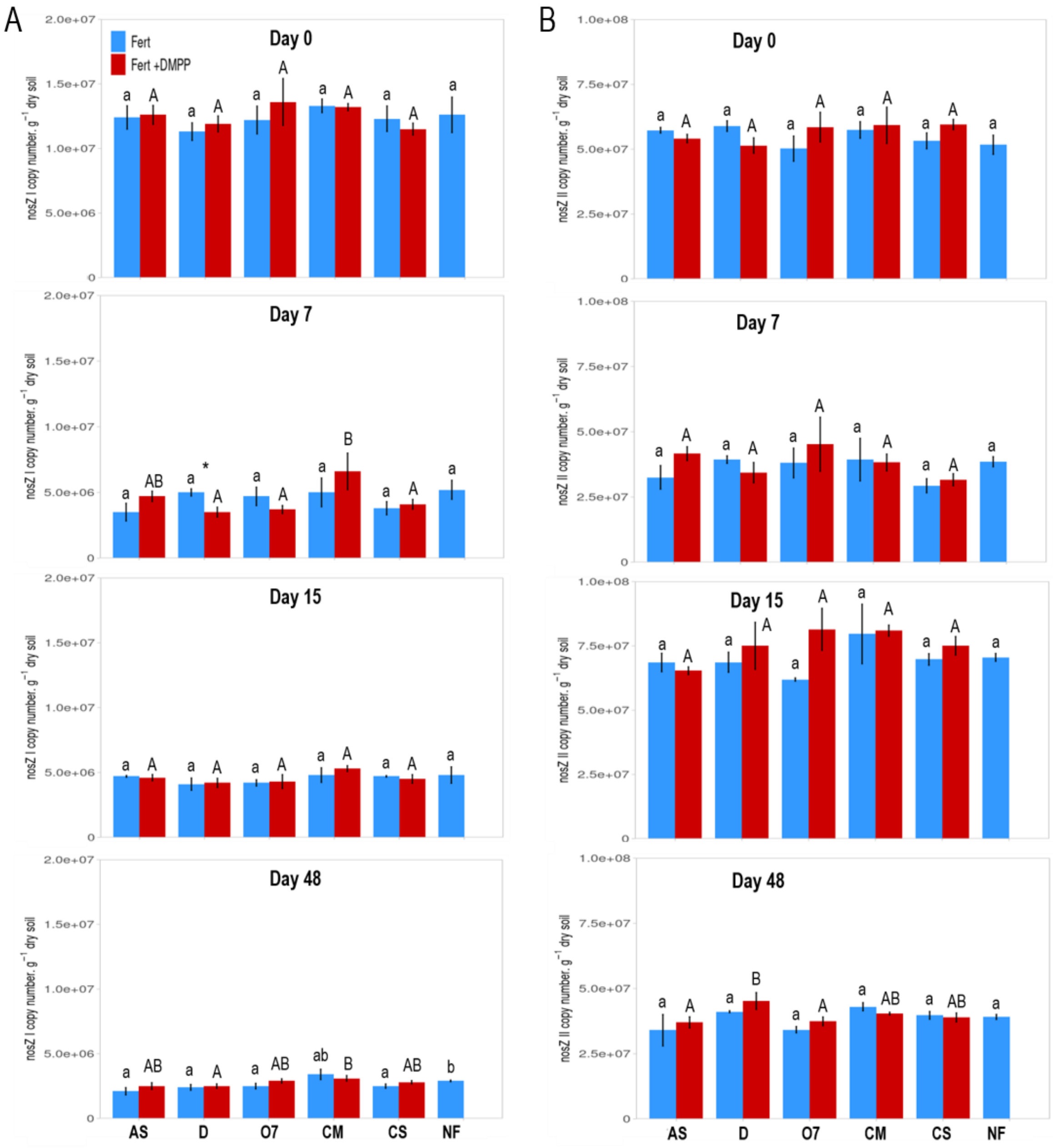


**Supplementary Fig. S8.** Abundance of total marker genes *nosZ I* and *nosZ II* determined at Day 0, 7, 15 and 48 after fertilization. Values represent mean ± SE (n = 4). Statistics as in Supplementary Fig. S3.

**Supplementary Table S7.** Pairwise comparison of bacterial community structure for total bacteria, the class of *Nitrospira* and the family of *Nitrosomonaceae* as affected by incubation time. Bold letters show significant differences at $p < 0.05$, underlined letters show statistical trends ($p = 0.05$-$0.1$).

| **Factors** | | | | | | | **Total bacteria** | | **Family *Nitrosomonaceae*** | | **Class N*itrospirae*** | |
|---|---|---|---|---|---|---|---|---|---|---|---|---|
| **time** | **fert** | **dmpp** | | **time** | **fert** | **dmpp** | **F value** | **p value** | **F value** | **p value** | **F value** | **p value** |
| | AS | NO | | | AS | NO | 1.259 | **0.035** | 1.685 | **0.035** | 0.207 | 0.968 |
| | O7 | NO | | | O7 | NO | 1.332 | **0.033** | 0.989 | 0.431 | 1.572 | 0.271 |
| | CM | NO | | | CM | NO | 1.625 | **0.041** | 2.062 | **0.041** | 0.935 | 0.494 |
| | D | NO | | | D | NO | 2.091 | **0.028** | 1.407 | **0.029** | 2.998 | **0.029** |
| | CS | NO | | | CS | NO | 2.136 | **0.024** | 1.312 | 0.109 | 3.583 | **0.024** |
| 7 | NF | NO | vs. | 48 | NF | NO | 1.321 | **0.033** | 1.098 | 0.362 | 0.899 | 0.490 |
| | AS | YES | | | AS | YES | 1.180 | **0.027** | 1.336 | 0.134 | 0.769 | 0.621 |
| | O7 | YES | | | O7 | YES | 1.298 | **0.019** | 0.913 | 0.649 | 0.405 | 0.906 |
| | CM | YES | | | CM | YES | 1.961 | **0.027** | 1.335 | <u>0.085</u> | 0.748 | 0.650 |
| | D | YES | | | D | YES | 1.902 | **0.037** | 1.978 | **0.037** | 2.745 | <u>0.064</u> |
| | CS | YES | | | CS | YES | 2.048 | **0.025** | 1.089 | 0.349 | 0.991 | 0.493 |

**Supplementary Table S8.** Pairwise comparison of bacterial community structure for total bacteria, the class of ***Nitrospira*** and the family of *Nitrosomonaceae* as affected by mineral (AS) vs organic fertilization. Bold letters show significant differences at p < 0.05, underlined letters show statistical trends (p = 0.05-0.1). D, liquid digestate; O7, Orgamine 7 pellets; CM, chicken manure; CS, cattle slurry (CS); NF, non-fertilized control.

| Factors | | | | | | Total bacteria | | Family *Nitrosomonaceae* | | Class *Nitrospirae* | |
|---|---|---|---|---|---|---|---|---|---|---|---|
| time | fert | dmpp | | fert | dmpp | F value | p value | F value | p value | F value | p value |
| 7 | AS | NO | vs. | O7 | NO | 1.244 | **0.021** | 1.346 | 0.115 | 0.997 | 0.453 |
| | | | | CM | NO | 1.600 | **0.036** | 1.489 | 0.063 | 0.970 | 0.470 |
| | | | | D | NO | 2.126 | **0.023** | 1.449 | **0.050** | 2.449 | 0.052 |
| | | | | CS | NO | 2.055 | **0.021** | 1.348 | 0.163 | 3.834 | **0.021** |
| | | | | NF | NO | 1.043 | **0.035** | 1.991 | **0.030** | 2.169 | 0.160 |
| | | YES | vs. | O7 | YES | 1.233 | **0.025** | 0.933 | 0.660 | 0.547 | 0.836 |
| | | | | CM | YES | 1.876 | **0.025** | 1.117 | 0.367 | 0.926 | 0.518 |
| | | | | D | YES | 2.083 | **0.036** | 1.665 | 0.089 | 0.732 | 0.616 |
| | | | | CS | YES | 2.163 | **0.033** | 0.840 | 0.640 | 1.369 | 0.346 |
| 48 | AS | NO | vs. | O7 | NO | 1.384 | **0.021** | 1.144 | 0.293 | 1.622 | 0.172 |
| | | | | CM | NO | 1.537 | **0.042** | 2.280 | **0.042** | 2.635 | 0.072 |
| | | | | D | NO | 1.460 | **0.033** | 1.021 | 0.542 | 0.977 | 0.587 |
| | | | | CS | NO | 2.163 | **0.027** | 1.445 | 0.102 | 1.081 | 0.484 |
| | | | | NF | NO | 1.321 | **0.026** | 1.806 | **0.026** | 1.544 | 0.119 |
| | | YES | vs. | O7 | YES | 1.116 | **0.027** | 0.888 | 0.668 | 0.514 | 0.809 |
| | | | | CM | YES | 1.539 | **0.035** | 1.103 | 0.329 | 0.675 | 0.704 |
| | | | | D | YES | 1.396 | **0.028** | 1.356 | 0.143 | 1.312 | 0.280 |
| | | | | CS | YES | 1.963 | **0.030** | 1.175 | 0.329 | 1.243 | 0.237 |

**Supplementary Table S9.** List of bacteria indicative for presence or absence of DMPP at T7 with a correlation strength cor > 0.45 as assessed by multipattern analysis. Table shows taxonomic classification and correlation strength

| | | **cor** | **p value** | **Phylum** | **Class** | **Order** | **Family** | **Genus** | **Species** | **relative** | **MeanCount** | **SumCount** |
|---|---|---|---|---|---|---|---|---|---|---|---|---|
| No DMPP | ZOTU9474 | 0.613 | 0.001 | Proteobacteria | Gammaproteobacteria | Burkholderiales | Nitrosomonadaceae | Nitrosospira | NA | 0.03533189 | 15.4545455 | 680 |
| | ZOTU14443 | 0.47 | 0.004 | Methylomirabilota | Methylomirabilia | Rokubacteriales | WX65 | WX65 | uncultured_bacterium | 0.01293771 | 5.65909091 | 249 |
| With DMPP | ZOTU7684 | 0.513 | 0.002 | Latescibacterota | Latescibacterota | Latescibacterota | Latescibacterota | Latescibacterota | NA | 0.00306556 | 1.34090909 | 59 |
| | ZOTU4244 | 0.474 | 0.004 | Planctomycetota | OM190 | OM190 | OM190 | OM190 | uncultured_bacterium | 0.01226224 | 5.36363636 | 236 |
| | ZOTU14032 | 0.464 | 0.003 | Acidobacteriota | Vicinamibacteria | Vicinamibacterales | Vicinamibacteraceae | Vicinamibacteraceae | uncultured_bacterium | 0.00904081 | 3.95454545 | 174 |
| | ZOTU13620 | 0.455 | 0.001 | Proteobacteria | Alphaproteobacteria | Rhizobiales | Xanthobacteraceae | Pseudolabrys | uncultured_bacterium | 0.02052367 | 8.97727273 | 395 |

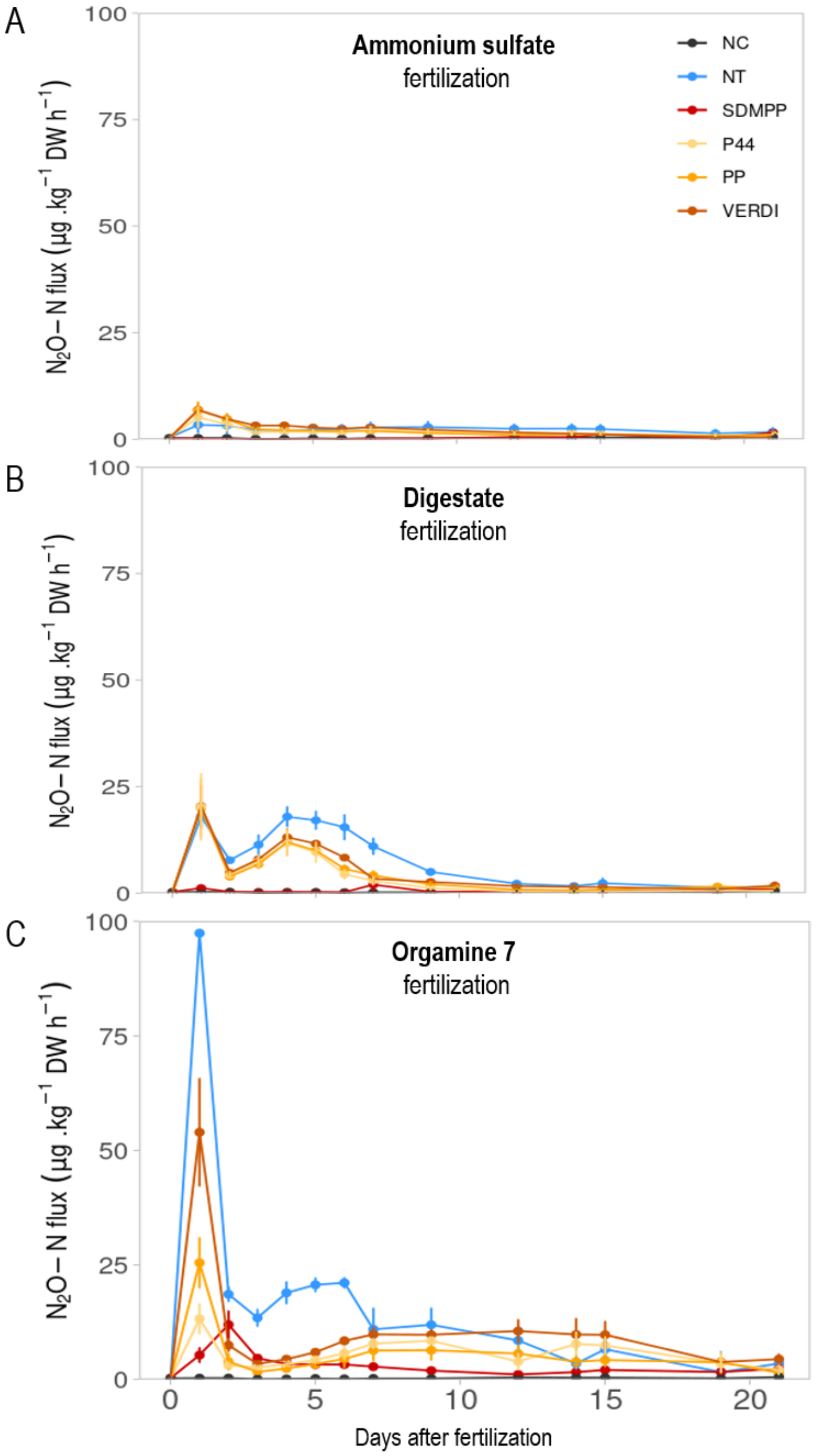


**Supplementary Fig. S9.** Daily $N_2O$ emission fluxes (EXP2) for 21-days incubation period. Incubation consisted in untreated and non-fertilized control soil (NC), fertilized not treated soil (NT), soil treated with DMPP (SDMPP), and soil collected from the rhizosphere of wheat cv. Persia 44 (P44), white mustard cv. Pole Position (PP) and cv. Verdi (VERDI). Soils were incubated with fertilizers (A) ammonium sulfate (AS), (B) liquid digestate (D) and (C) Orgamine 7 (O7). For each fertilizer, letters represent significant differences between soil treatments analyzed by Duncan's test (P<0.05) of each soil treatment with fertilizers AS, D and O7.
